# High-throughput identification of changes in cellular localization for RNA-binding proteins during adenovirus infection reveals viral harnessing of nuclear speckles

**DOI:** 10.64898/2026.11.16.765444

**Authors:** Amber R. N. Abbott, Tomer Lapidot, Tanner M. Tessier, Songtao Ye, Holly Chan, Edwin Halko, Giorgia Piserchia, Danfeng Cai, Ophir Shalem, Alexander M. Price, Matthew D. Weitzman

## Abstract

Adenovirus (AdV) is a human DNA virus reliant on host RNA processing machinery. While research into AdV has yielded landmark discoveries in RNA biology, studies investigating the scope of AdV dependency on specific host RNA-binding proteins (RBPs) are lacking. Here we performed optical pooled screening on a cell library where hundreds of RBPs are endogenously tagged. Pooled imaging followed by *in situ* barcode sequencing revealed a broad view of virus-induced host RBP relocalization patterns, including those recruited to viral replication compartments (VRCs). A targeted siRNA screen of relocalized RBPs demonstrated a requirement for the nuclear speckle (NS) core protein SON during virus infection. Depletion of SON resulted in reduced viral gene expression, impaired NS protein recruitment to early-stage VRCs, and impaired mobility of SRRM2 during infection. These findings suggest a crucial role of SON and NS functions in regulating viral gene expression and demonstrate the power of pooled tagging coupled to optical screening in investigating virus-host interactions.

## INTRODUCTION

Adenoviruses (AdV) are highly prevalent vertebrate pathogens that can induce a wide range of disease states^1^. While infections in immunocompetent individuals are typically self-limiting, immunocompromised individuals or those with pre-existing conditions are at risk of more severe disease^2,3^. Together with the lack of antivirals or vaccines approved for civilian use, these viruses represent a threat to public health^4,5^. Additionally, these viruses are a subject of great interest due to their capacity to serve as tools for vaccine and therapeutic applications^6–8^. AdV-based vaccines have been generated against numerous pathogens, including Ebola^9^ and SARS-CoV-2^10,11^. The clinical relevance of AdV, due to both the disease burden they induce as pathogens themselves, as well as the therapeutic potential they possess as vectors to combat other diseases, demands a thorough understanding of the interplay between these viruses and the host cells they infect.

Research into AdV biology has resulted in numerous seminal discoveries in virology and cell biology^12^. Since AdV is a nuclear-replicating DNA virus that depends on host cell machinery to facilitate RNA processing, many of these discoveries have significantly advanced our understanding of cellular RNA biology. The investigation of AdV transcripts led to fundamental insights into RNA capping^13^ and alternative polyadenylation^14^, as well as the discovery of splicing^15^. These landmark discoveries greatly expanded our understanding of mechanisms utilized by AdV to facilitate expression and processing of viral transcripts. However, many studies investigating AdV RNA biology have been limited in scope, and therefore the extent of AdV dependency on specific host RNA-binding proteins (RBPs) is not fully appreciated. We aimed to address this question by employing unbiased screening approaches that utilize pooled gene tagging to identify which RBPs are co-opted for viral RNA processing on a broader scale.

Until recently, the analysis of changes in protein localization patterns relied on methods that were relatively limited in scale. These include antibody-based techniques and the endogenous or exogenous tagging of individual proteins of interest. However, recent scaling of gene-tagging technologies has enabled the generation of pooled cell libraries, where each cell carries a unique endogenously tagged protein^16–20^. Here we used Scalable POoled Targeting with a LIgandable Tag at Endogenous Sites (SPOTLITES) where homology-independent repair is used to insert a generic synthetic exon encoding HaloTag into introns of endogenous genes^20,21^. The resulting cell libraries can be analyzed with optical pooled screening approaches which employ *in situ* barcode sequencing to pair the identity of the protein of interest with the corresponding localization pattern^22^. Optical pooled screening of cells in other contexts has been used to identify regulators of viral infections for both Sendai^23^ and Ebola^24^ viruses. These studies employed image-based CRISPR knockout screens to identify cellular proteins impacting infection. However, the primary focus of these experiments was to identify the impact of genetic perturbations on viral protein and/or RNA localization patterns, and therefore the number of host proteins included in the respective image analyses was limited. We reasoned that optical pooled screening of a SPOTLITES library following infection would provide a broad view of virus-induced host protein relocalization patterns, including those recruited to viral structures. We further reasoned that any host proteins which relocalized to viral structures may represent proteins co-opted to promote infection, and these would therefore be ideal candidates for secondary functional screening.

AdV infection induces the formation of membrane-less nuclear compartments termed viral replication compartments (VRCs). In addition to facilitating viral genome replication, these compartments are thought to organize other key viral processes spatially within the infected nucleus, including viral gene expression^25–27^. AdV is known to recruit cellular factors to VRCs to aid these processes^28–31^. Surrounding the sites of viral DNA replication are the sites of viral RNA transcription^25,32^. Multiple cellular RBPs have been shown to localize in this ring-like pattern, including RNA Pol II^33^, multiple m^6^A-interacting proteins^34^, and small nuclear ribonucleoprotein (snRNP) splicing factors^25,33,35,36^. We reasoned that the localization pattern of AdV VRCs would be an ideal metric for comparison for an image-based screen, with the goal of determining host proteins recruited to promote viral processes, including viral RNA processing.

We infected a cell library containing 778 RBPs endogenously tagged with HaloTag, followed by image-based phenotyping in combination with *in situ* sequencing of the corresponding sgRNA^20^. We then employed a deep learning algorithm to identify AdV-induced RBP relocalization patterns, including those which co-localized with VRCs. A secondary siRNA screen of RBPs recruited to VRCs revealed a requirement for the nuclear speckle (NS) core protein SON during AdV infection. We determined that SON is required for efficient viral gene expression, and that SON knockdown impairs recruitment of other NS proteins to sites of viral RNA synthesis. Live-cell imaging revealed that SON also impacts both the initial formation of VRCs as well as the proximity of VRCs and NS as infection progresses. Our findings support a model where SON mediates AdV RNA expression by facilitating the recruitment of speckles to nascent VRCs as well as the dynamic exchange of NS proteins with the nucleoplasm. We additionally demonstrate the power of this methodology for investigating virus-host interactions.

## RESULTS

### Identification of virus-induced relocalization patterns by high-throughput imaging of tagged RBPs

We recently generated a cell library by transducing an optimized lentiviral tagging library containing 1407 sgRNAs covering 778 genes into HeLa-M cells. These cells were also designed to express G3BP1 endogenously tagged with mClover3 to serve as a marker for stress granule formation. Since AdV does not induce stress granules^37^, we instead utilized this channel as a cytoplasmic marker to enable cell segmentation in the subsequent analysis. We infected the cell library with wild-type (WT) HAdV-C5 at a multiplicity of infection (MOI) of 10 (**Figure 1A**). At 23.5 hours post-infection (hpi), the fluorescent JFX549 HaloTag ligand was added for 15 minutes (min) followed by 3 washes, and the cells were then fixed for imaging at 24 hpi. A second plate infected in parallel was stained with an antibody recognizing the AdV DNA-binding protein (DBP) to confirm that essentially every cell was infected (**Figure S1A**). Following fixation, cells in the experimental plate were stained with DAPI and then imaged to ascertain the localization pattern of each cellular RBP. This was achieved as previously described by automated imaging of three channels consisting of each unique RBP tagged with HaloTag, cytoplasmic G3BP1 tagged with mClover3, and the nucleus stained with DAPI^20^. The entire area of each well in a 6-well plate was imaged. The identity of each tagged RBP within its corresponding cell was then ascertained by *in situ* sequencing following an established protocol^22^. Briefly, the sgRNA sequence was reverse-transcribed and amplified by rolling circle amplification (RCA) to generate a localized DNA sequence that was determined by consecutive rounds of sequencing by synthesis (SBS) and imaging. Following 14 rounds of SBS, the RBP phenotypes for each cell were linked to their corresponding genotype, resulting in a total of 1,211,041 identified cells. We found that approximately half of the RBPs were tagged in a single intron, with another quarter represented by tagging in two different introns, and a final quarter represented by three different introns (**Figure S1B**). We identified multiple diverse subcellular localization patterns, and generated “cell albums” for each tagged protein (**Figure 1B**). We observed that the majority of RBPs displayed either a diffuse, nuclear, or cytosolic subcellular distribution (**Figure 1C**). Strikingly, many nuclear RBPs demonstrated a pattern of clustered rings resembling that of VRC formation (**Figure 1D**). We selected a few proteins for validation by immunostaining, and confirmed that they indeed relocalize to surround VRCs during infection of the parental HeLa cells expressing endogenously tagged G3BP1 (**Figure S1C**). Collectively, these data demonstrate our ability to define protein localization patterns following viral infection and optical pooled screening of a SPOTLITES cell library, and suggest that infection likely induces recruitment of many nuclear RBPs to VRCs.

**Figure 1:**
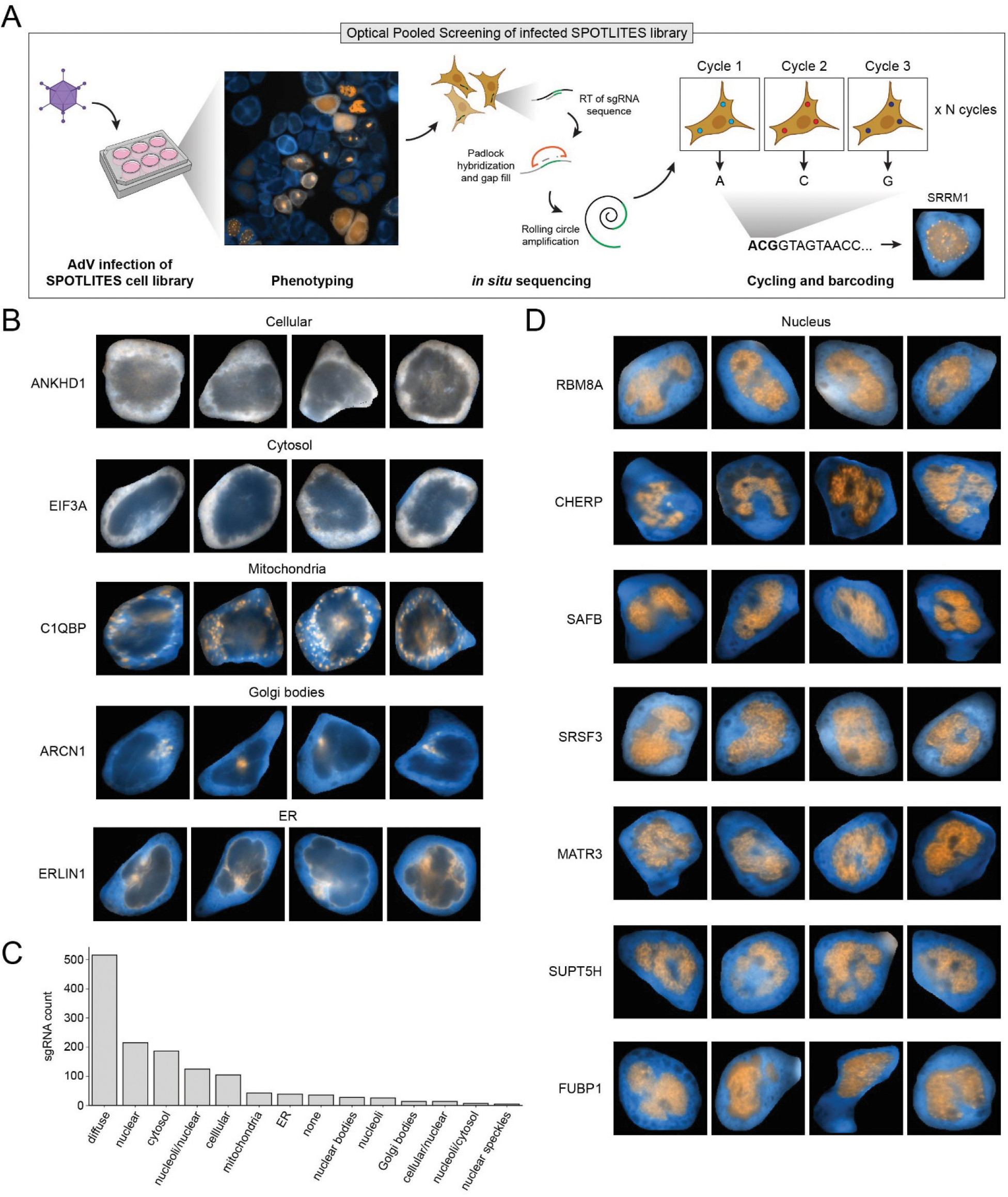
Optical pooled screening enables identification of virus-induced RBP relocalization patterns. A) Schematic of experimental workflow for the screen. B) Image albums of proteins demonstrating various localization patterns in infected cells. C) Number of sgRNAs corresponding to each identified localization pattern. D) Image albums of nuclear proteins demonstrating a pattern similar to that of viral replication compartments.

### AdV infection induces relocalization of nuclear RBPs to viral replication compartments

To determine virus-induced changes to RBP localization patterns, we applied a deep learning approach to compare our dataset with another dataset of the same library which was uninfected. We first compared the sgRNA counts between the infected and uninfected datasets to ensure reproducibility and found a high correlation between the two (**Figure S2A**). We then trained a variational autoencoder (VAE)^38^ to recognize VRCs. We generated a training dataset by infecting the parental HeLa cells with AdV for either 16 or 24 hpi and staining for DBP to demark VRCs. We then performed cellular segmentation on the G3BP1 channel using Cellpose, resulting in 30,000 cropped single-cell images. In addition to this viral phenotype, the VAE was trained on a semi-manually curated dataset representing 12 distinct cellular localization patterns (with a total of 320,184 images across all 13 patterns). These patterns include the cytoplasm, nucleus, nucleoli, endoplasmic reticulum (ER), and mitochondria (**Figure S2B)**. This workflow enabled it to distinguish both normal and infection-associated phenotypes within a shared latent space. After training, the VAE was used to infer localization patterns across the remaining cells in both datasets. The resulting latent representations were projected into a UMAP embedding to visualize relationships between localization phenotypes (**Figure 2A**). Representative single-cell images from each cluster demonstrated that the model accurately distinguished distinct localization patterns for each tagged protein, including proteins localizing to viral structures.

**Figure 2:**
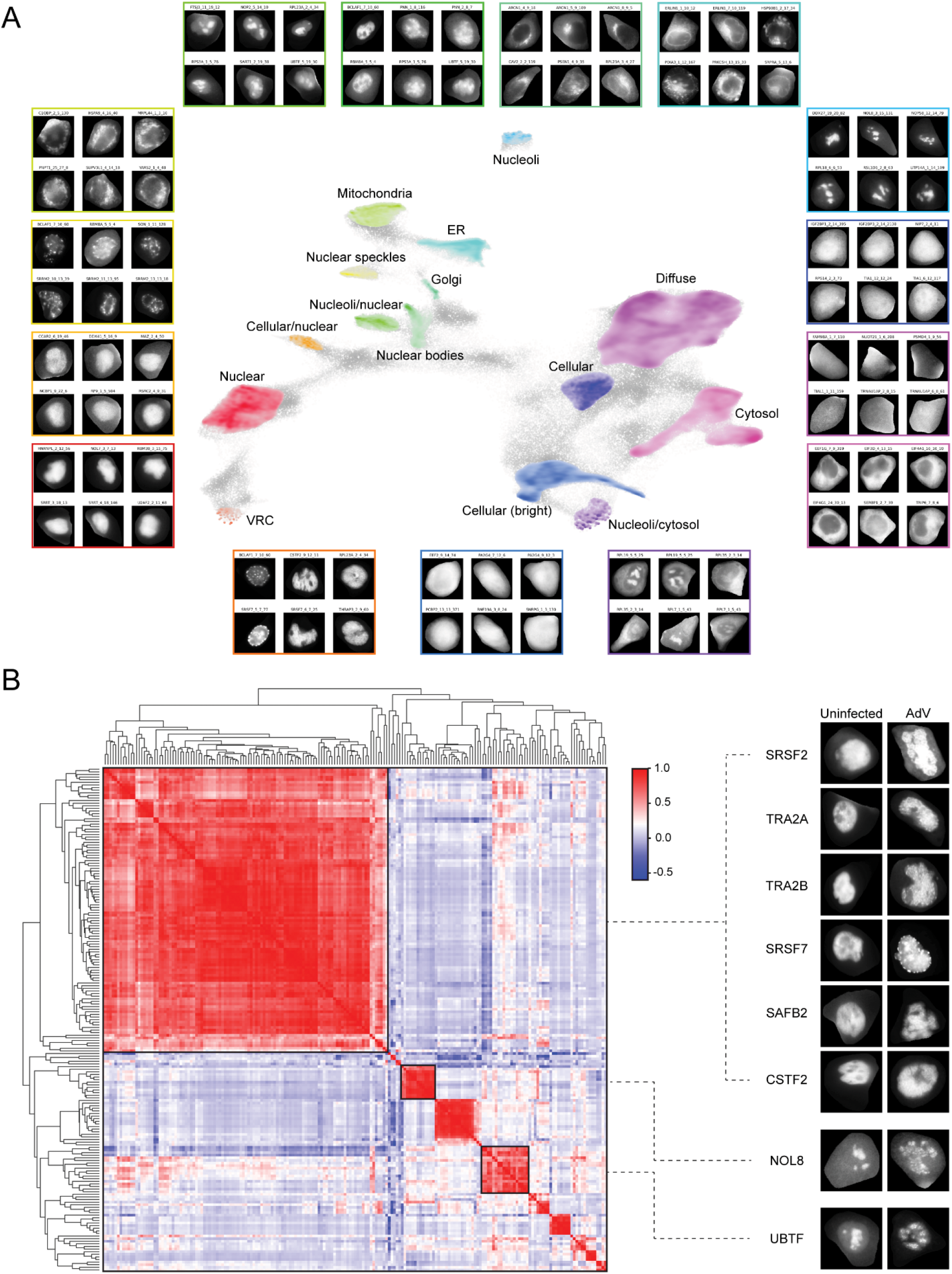
AdV infection induces relocalization of nuclear RBPs to viral replication compartments. A) UMAP displaying clusters of cells with RBPs demonstrating the indicated localization pattern. Representative images of cells within each corresponding cluster are shown. B) Heatmap of cosine similarity between movement of cell clusters in the latent space due to phenotypic change induced by AdV infection. Single-cell images are shown from each group that represent relocalization to a VRC-like pattern.

To quantify changes in localization patterns between normal and infected states for each sgRNA, silhouette scores were calculated using the 64-dimensional latent representations generated by the VAE. Specifically, silhouette scores were computed between control and infected cell populations to measure the magnitude of phenotypic separation induced by infection. To determine the direction and type of localization change associated with each sgRNA, the cosine similarity of the shift vector in latent space was calculated between the centroids of the normal and infected cell populations. Together, this analysis enabled quantification of both the magnitude of phenotypic change (via silhouette score), as well as the nature of the localization transition (via angular displacement in latent space) on a per-sgRNA basis. This analysis revealed that relocation from the nucleus to VRCs was the predominant relocalization pattern with the largest number of cellular proteins (**Figure 2B**). Additional patterns included relocalization of nucleolar proteins to VRCs (as exemplified by NOL8), as well as accumulation at VRCs for proteins normally present in both nucleoli and the nucleoplasm (as exemplified by UBTF). These data demonstrate that AdV infection induces relocalization of cellular RBPs on a broad scale, including a striking number of nuclear RBPs recruited to replication compartments.

### A targeted siRNA screen of relocalized RBPs reveals proteins required for AdV infection

Our next goal was to determine whether this virus-induced rearrangement of RBPs may reflect a viral strategy to promote infection. To identify which relocalized RBPs are required for AdV infection, we performed a secondary siRNA screen targeting genes with the highest silhouette scores from each of the RBP groups in **Figure 2B**. We tested a total of 80 targets, and included POLR2A (the catalytic core subunit of RNA Pol II) as a positive control. This approach allowed us to focus on proteins recruited to VRCs, but also gain additional information regarding the relevance of proteins which demonstrate alternative virus-induced relocalization patterns. As a readout for viral progeny, we utilized the previously published reporter virus AdNTE, which was designed to express NanoLuc (NLuc) luciferase upstream of the E1A gene^39^. NLuc is linked to E1A by a porcine teschovirus 2A (P2A) self-cleaving peptide sequence in order to uncouple reporter activity from E1A function. Not only is this virus a useful tool to analyze changes in early gene expression, but bioluminescent quantification of infectious viral progeny using AdNTE has been shown to be as accurate as quantification by standard plaque assay^39^. We performed the screen in A549 cells in a 96-well plate format (**Figure 3A**). Cells were transfected with the siRNA library for 24 h before infection with AdNTE, and then samples were harvested for quantification at 24 or 48 hpi. New A549 cells were infected with viral supernatants, and luciferase activity analyzed at 6 hpi according to the established protocol^39^. Relative light unit (RLU) values for each sample were compared to those of their respective internal nontargeting negative control. The screen was performed twice to ensure reproducibility. Results at 24 and 48 hpi yielded *R*^2^ values of 0.5554 and 0.8384, respectively, indicating that the findings were consistent across experiments (**Figure 3B**). To ensure that siRNA treatment alone did not impact cell viability, we transfected cells with the same library for 3 days before assessing relative cell viability with a luminescent CellTiter-Glo 2.0 assay (**Figure S3A**). Results demonstrated only one siRNA with a substantial impact on RLU values (corresponding to NXF1), so this target was excluded from further analysis (**Figure S3B-C**).

**Figure 3:**
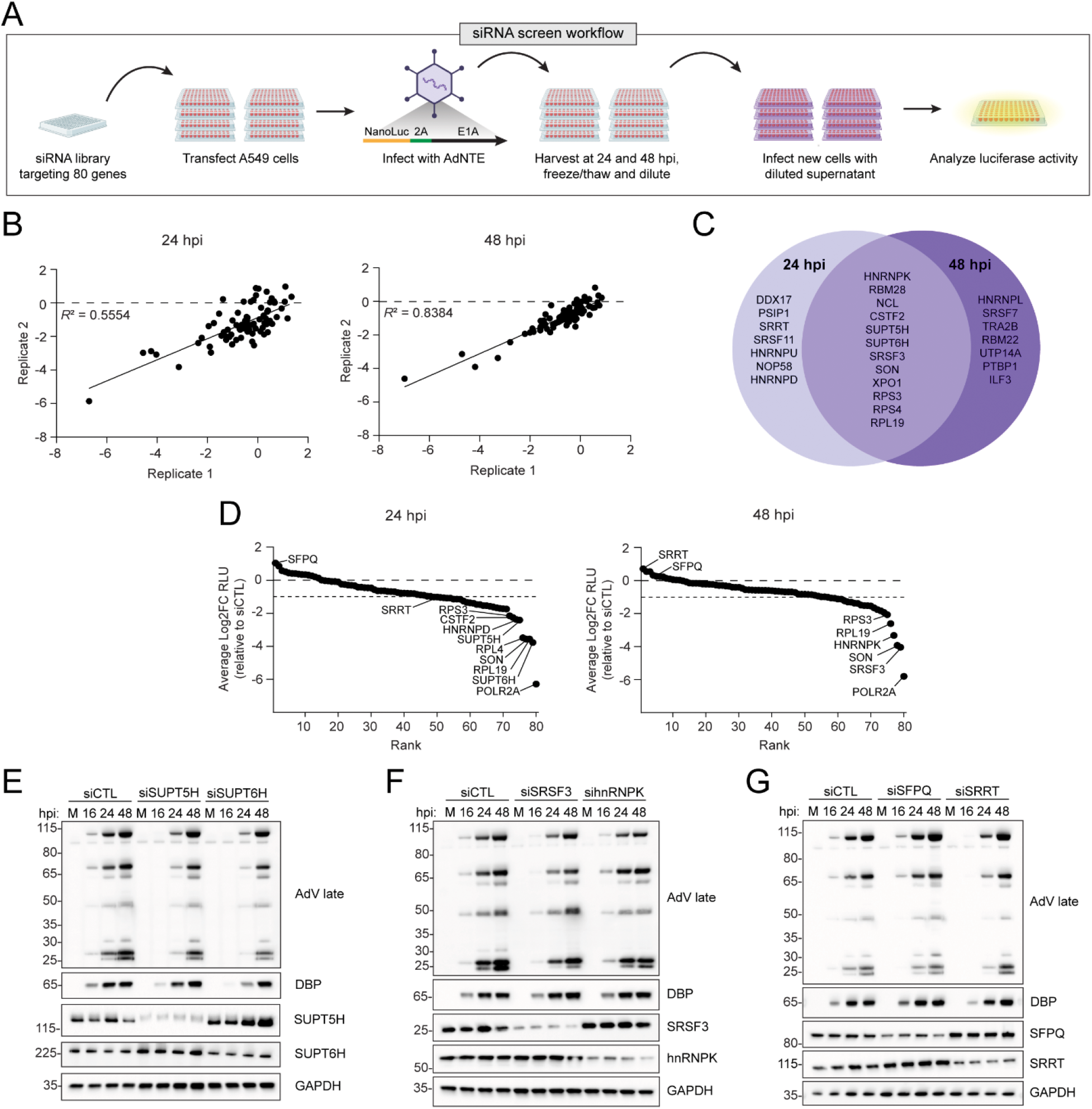
A targeted siRNA screen of relocalized RBPs reveals proteins required for AdV infection. A) Schematic for siRNA screen of genes required for AdV infection. B) Reproducibility of the screen was analyzed by comparing the Log2FC of the blank-adjusted RLU values relative to the nontargeting siCTL for each replicate. Coefficient of determination is displayed for each timepoint. C) Venn diagram depicting genes which induced a Log2FC<-1 RLU relative to the nontargeting siCTL for both replicates at one or both timepoints when silenced. D) Rank plots for each timepoint of the siRNA screen. Genes were ranked by the Log2FC blank-adjusted RLU values relative to the nontargeting siCTL, averaged for each replicate. Genes which induced a Log2FC<-2 are indicated, along with SFPQ and SRRT. E) Immunoblot depicting early (DBP) and late viral protein levels over a timecourse following knockdown of SUPT5H or SUPT6H for 24 h. F) Immunoblot depicting early (DBP) and late viral protein levels over a timecourse following knockdown of SRSF3 or hnRNPK for 24 h. G) Immunoblot depicting early (DBP) and late viral protein levels over a timecourse following knockdown of SFPQ or SRRT for 24 h.

We analyzed the impact of each siRNA on AdNTE luciferase activity by determining the Log2FC in RLU values relative to the nontargeting control. The results revealed 12 genes that when silenced induce a Log2FC < −1 at both timepoints, 7 genes with a similar impact only at 24 hpi, and 7 with an impact only at 48 hpi (**Figure 3C**). Ranking the effects of each gene revealed that knockdown of the nuclear speckle core protein SON had the third-most pronounced impact on viral infection at 24 hpi, and the second-most pronounced impact at 48 hpi (**Figure 3D, S3D**). Other silenced genes with pronounced impacts at 24 hpi include transcription elongation factors SUPT5H and SUPT6H, the ribosomal proteins RPS3, RPL4, and RPL19, heterogenous nuclear ribonucleoprotein D (hnRNPD), and cleavage stimulation factor subunit 2 (CSTF2). Genes with the most pronounced impact at 48 hpi when silenced include SRSF3 and hnRNPK. At 24 hpi, SFPQ was the only gene that resulted in a substantial positive impact on infection. At 48 hpi, no gene induced a positive impact above the cutoff, but SRRT knockdown induced the strongest positive effect. We validated several of these findings by analyzing protein production over a time-course of WT HAdV-C5 infection. Knockdown of SUPT5H and SUPT6H each resulted in lower levels of viral early (DBP) and late proteins at 16 and 24 hpi, with less of an impact at 48 hpi, reflecting the findings from the screen (**Figure 3E**). Knockdown of SRSF3 and hnRNPK had negligible impact on early protein levels at any timepoint but had a more pronounced impact on late proteins at 48 hpi (**Figure 3F**). Knockdown of SFPQ induced a mild increase in late proteins at 24 and 48 hpi, whereas knockdown of SRRT induced a stronger increase at 48 hpi (**Figure 3G**). These results validate our siRNA screening approach, and demonstrate we can detect impacts of proteins which exert proviral or antiviral effects at different stages of infection. Since knockdown of SON induced a striking decrease in luciferase activity at both timepoints, we chose this protein to investigate further functions during AdV infection.

### SON is required for efficient adenoviral gene expression

SON is a core component of nuclear speckles, which are membrane-less condensates of RBPs and RNAs considered to exhibit properties of phase-separated bodies^40–43^. Previous efforts to determine the components of speckles have demonstrated these bodies are enriched in pre-mRNA splicing factors, along with factors involved in transcription, mRNA processing and export, and messenger ribonucleoprotein (mRNP) packaging^44–46^. While speckles have been historically viewed as storage sites for splicing factors^47^, a growing number of studies have shown that speckles are associated with active genes^48–53^, thereby informing the current perspective that speckles are active hubs promoting gene expression. Speckles have been ascribed various roles during viral infections^54–62^. However, while SON has been shown to be required for infection of certain viruses including influenza^56,63,64^, human Hepatitis B Virus (HBV)^65^, and HIV^60^, the role of this protein during AdV infection has not been investigated.

We first tested the impact of knocking down SON, either individually or together with the other core protein SRRM2, on specific stages of the viral replication cycle. These two proteins together are required for speckle structure, and their combined depletion has been shown to result in speckle dissolution^66^. We reasoned that this method would allow us to distinguish between the impact of SON and that of the speckles as a whole, should there be any viral process only affected by the double knockdown. To validate the impact on progeny from the screen as determined by luciferase activity, we knocked down SON and/or SRRM2 in A549 cells for 24 h before infection, then harvested cells at 24 and 48 hpi and performed a classic plaque assay. The individual knockdown of SON as well as the double knockdown resulted in a significant decrease in the number of plaques compared to control siRNA at each timepoint (approximately 3-fold), while there was no impact of the individual SRRM2 knockdown (**Figure 4A**). We observed a similar impact on viral genome amplification at the same timepoints as determined by qPCR (**Figure 4B**). To assess the impact on the production of viral proteins, we harvested lysates at 16 and 24 hpi following SON and/or SRRM2 knockdown, and analyzed early and late protein levels by immunoblot analysis (**Figure 4C**). Results show a reduction in both early viral proteins (DBP and E4) and late structural proteins in response to the single knockdown of SON or the double knockdown. Intriguingly, this result was not observed for the E1A proteins, which were minimally affected. To determine whether these findings reflect an effect on viral gene expression, we performed reverse transcription coupled with quantitative PCR (RT-qPCR) to measure viral mRNA levels at 18 or 24 hpi (**Figure 4D**). Results show a significant decrease in expression of early and late genes following individual SON knockdown or the double knockdown, with the exception of E1A, which is the first viral gene to be expressed. At both timepoints, any changes in E1A expression were less than 2-fold, although there was a more pronounced negative impact of the double knockdown than either individual knockdown at 16 hpi. However, at 24 hpi E1A expression was significantly increased following knockdown of SON relative to the control, with a slight negative impact of the SRRM2 knockdown, and no effect of the double knockdown. We analyzed the splicing efficiency of early and late genes by determining the ratio of the spliced and unspliced transcript levels (**Figure S4**). The splicing efficiencies of E1A and the tripartite leader (TPL) of the Major Late Promoter (MLP) common to nearly all late genes were similar to results of their respective transcript levels. Given the significant impact of SON depletion on viral gene expression, we hypothesized that this protein may be required to facilitate transcription of viral genes. We therefore analyzed nascent RNA transcription with a 4-thiouridine (4SU) labeling approach. Following 24 h of SON knockdown, A549 cells were infected with AdV for 23 h, then pulsed with 4SU for 1 h before harvesting at 24 hpi. Labeled RNA was enriched and analyzed by RT-qPCR. We analyzed unspliced regions near the E1A and E4 regions, as well as within the TPL (**Figure 4E**). Consistent with the total viral mRNA levels, the results indicate significantly increased relative E1A transcription, in contrast to decreased relative E4 and TPL transcription. Collectively, these findings suggest that SON is required for efficient expression of viral genes following expression of E1A. Since the double knockdown of SON/SRRM2 showed no further decreases in viral processes we chose to focus on SON-specific RNA processing defects, as opposed to those that could be mediated by the NS structure as a whole.

**Figure 4:**
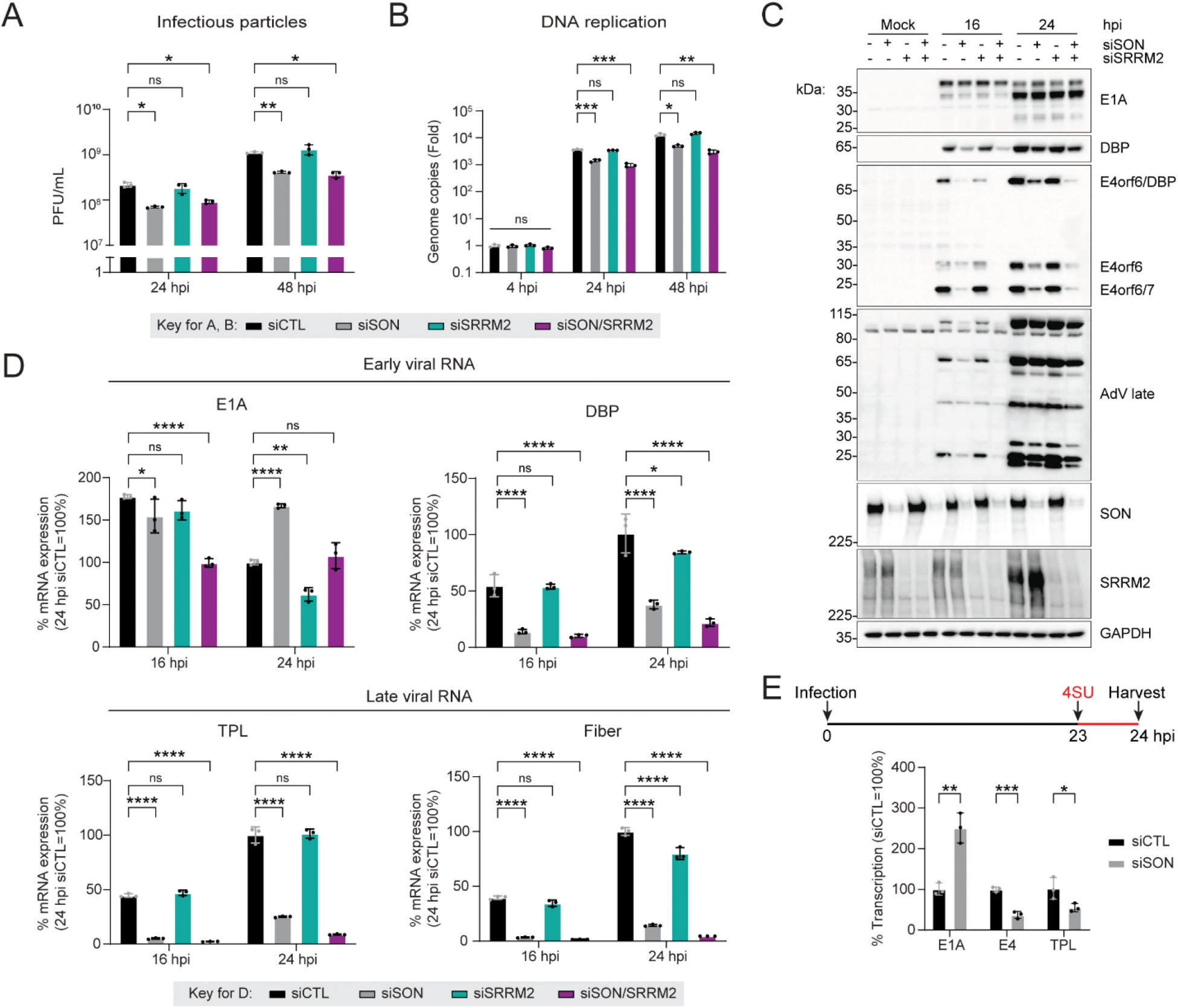
SON is required for efficient adenoviral gene expression. A) Infectious particle production was determined by plaque assay following knockdown of speckle core proteins SON and/or SRRM2 in A549 cells. B) Viral DNA replication was determined by qPCR at different times after infecting A549 cells depleted of SON and/or SRRM2 by siRNA. C) Levels of viral late proteins or early (E1A, DBP, E4orf6) proteins at 16 and 24 hpi following SON and/or SRRM2 knockdown for 24 h. D) Viral RNA expression was measured by RT-qPCR after SON and/or SRRM2 knockdown. E) Nascent transcription was analyzed by labeling RNA with 4sU for 1 h in A549 cells following knockdown of SON. Cells were harvested at 24 hpi, and 4sU-labeled RNA was extracted for analysis by RT-qPCR. Relative transcription rates of two early genes (E1A and E4) and the tripartite leader (TPL) found in all late transcripts. Significance was determined by two-way ANOVA with Dunnett’s multiple comparisons test (A, B, D) or unpaired two-tailed Student’s *T*-test (E). *p≤0.05, **p≤0.01, ***p≤0.001, ****p≤0.0001, ns=not significant. All graphs represent mean +/- standard deviation.

### SON is required for SRRM2 relocalization to sites of AdV RNA transcription

SON has been proposed to act as a scaffold for other NS components^67^. We hypothesized that loss of this protein would impair the ability of RNA-processing factors to access and mediate expression of viral RNA. Supporting this theory, AdV has been previously shown to induce reorganization of known NS components into rings around early replication compartments at sites of transcription, followed by formation of enlarged NS at the nuclear periphery later in infection^35^. We first examined images from the optical pooled screen and confirmed that both SON and SRRM2 exhibit this enlarged NS pattern during infection (**Figure 5A**). We then examined other NS proteins detected in the screen and found that many also demonstrate a similar pattern. We validated this phenotype of SON by immunofluorescence in A549 cells infected for 20 h (**Figure S5A**). This timepoint displays a heterogeneous population of cells demonstrating different stages of infection, allowing investigation of SON localization within cells during early, middle, and late stages of infection. In the early stage, SON forms striking rings around nascent VRCs marked by DBP. We corroborated this finding in HeLa cells expressing endogenous SRRM2 tagged with mScarlet, and confirmed that the two core proteins colocalize (**Figure 5B**). By the late stage of infection, when DBP staining is present throughout the nucleus, each protein exhibits the enlarged speckle phenotype detected in the optical pooled screen. Since growing evidence suggests that NS are associated with actively-transcribing genes, we next tested whether recruitment of SRRM2 to rings around early-stage VRCs is dependent on transcription. Immunostaining with an antibody recognizing phosphoserine 2 on the RNA Pol II C-terminal domain (Pol II p-Ser2) as a marker of actively transcribing polymerase confirmed that active Pol II surrounds DBP-marked VRCs, and colocalizes with SRRM2 rings (**Figure 5C**). Concurrently, we infected HeLa mScarlet-SRRM2 cells for 18 h, and subsequently treated them with transcription inhibitor DRB for 2 h. In cells treated with DRB, SRRM2 no longer demonstrates the ring pattern. We also found that this effect is reversible following removal of the inhibitor, with both phosphorylated Pol II and SRRM2 reforming ring-like structures. These findings demonstrate that NS core proteins are recruited to nascent VRCs in a transcription-dependent manner, and that they transition into larger speckles as infection progresses and VRC morphology changes.

**Figure 5:**
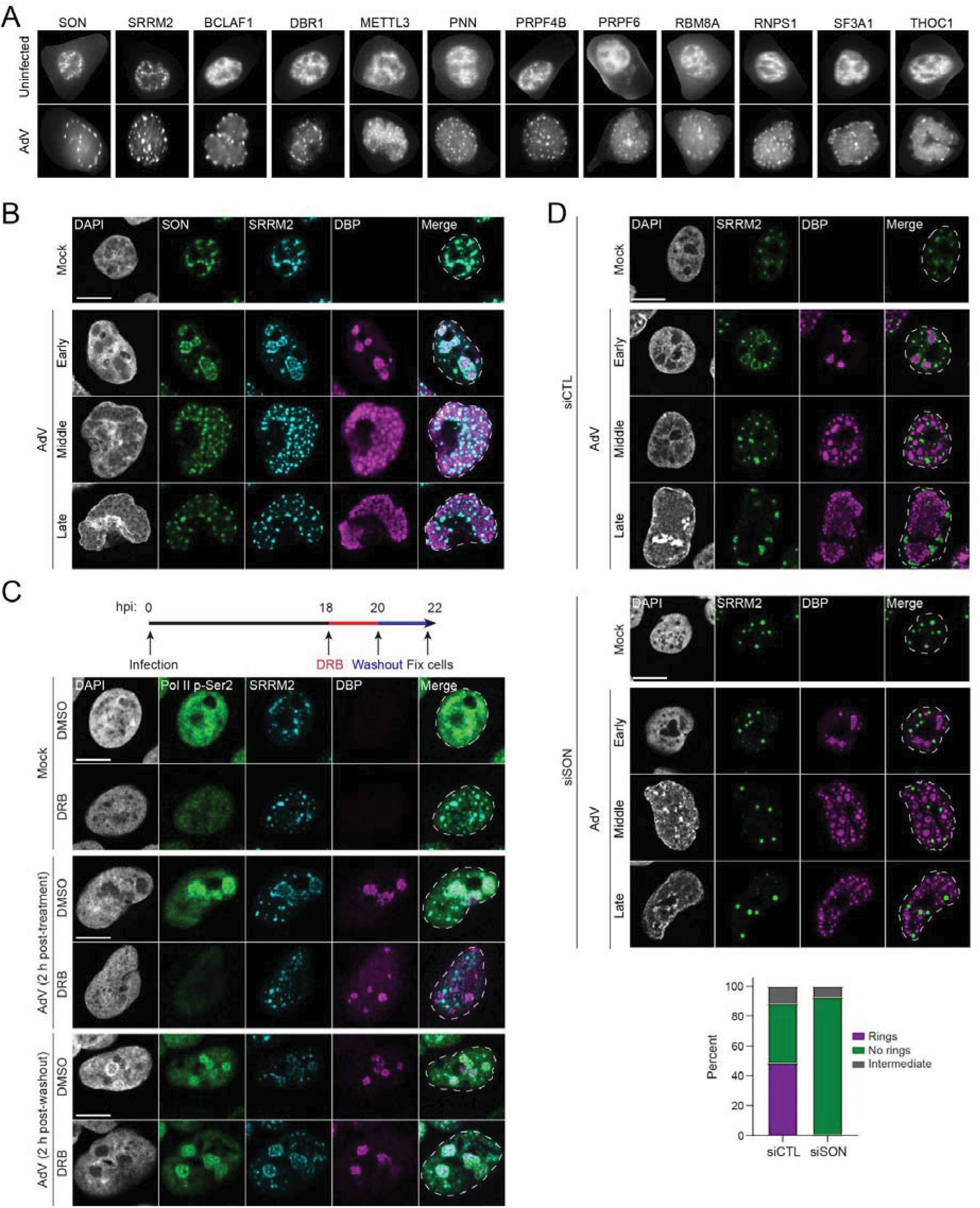
SON knockdown prevents SRRM2 from relocalizing to sites of AdV RNA transcription. A) Images of proteins enriched in NS in infected and uninfected cells from the optical pooled screen, including core proteins SON and SRRM2. B) Confocal images depicting SON localization in HeLa mScarlet3-SRRM2 cells following infection with AdV for 20 h. C) Confocal images of HeLa mScarlet3-SRRM2 cells following infection with AdV for 18 h, then treatment with 60 mM DRB for 2 h. At 20 hpi, the inhibitor was washed out for 2 h and cells fixed at 22 hpi. Pol II p-Ser2 (green) denotes actively transcribing RNA Polymerase II phosphorylated on serine 2 of CTD. D) Confocal images showing the pattern of SRRM2 in A549 cells relative to viral replication centers following knockdown of SON. All cells were fixed at 20 hpi, and representative cells in different stages of infection are shown (top). Quantification of cells in the early stage of infection which demonstrate the formation of rings around VRCs (bottom). 154 (siCTL) and 163 (siSON) total cells were quantified. B-D) DBP (magenta) is the viral DNA binding protein that marks sites of viral replication centers. The nuclear periphery is shown by a dotted white line as assessed by DAPI staining. Scale bar = 10 µm.

We next tested whether this recruitment of SRRM2 to sites of viral transcription depends on the presence of SON. We knocked down SON before infection for 20 h, then performed immunostaining for SRRM2 (**Figure 5D, S5B**). Similar to previous reports, loss of SON results in larger, more rounded speckles in uninfected cells^66–69^. Intriguingly, this SON deficient pattern is maintained throughout infection, and ring formation around VRCs no longer occurs. Quantification of cells in the early stage revealed ∼50% demonstrate these rings for the non-targeting siRNA control, whereas < 1% of cells demonstrate rings in the absence of SON (**Figure 5D**). We confirmed that this result is specific to SON by testing the impact of knocking down SRRM2, which does not impact ring formation of SON (**Figure S5C**). Collectively these results suggest that the presence of SON is required to allow for recruitment of the other NS core proteins to actively-transcribing viral RNA.

### SON is required for efficient dynamic exchange of NS proteins during infection and impacts VRC formation

We next addressed whether loss of SON impacts the ability of any other RBPs required for infection to relocalize to sites of viral RNA. We turned to our results of the siRNA screen, and tested whether proteins important for infection (**Figure 3C**) also showed reduced relocalization upon SON knockdown. We identified SRSF11, PSIP1, DDX17, SRSF3, and hnRNPD as proteins with decreased recruitment to VRCs following SON knockdown, as determined by the proportion of infected A549 cells in the early stage which exhibit the ring phenotype (**Figure 6A**). Intriguingly, these proteins also partially colocalized with SRRM2 in the HeLa mScarlet-SRRM2 cells (**Figure S6A**). These findings indicate that multiple proteins that are both enriched in NS and relevant to infection require the presence of SON in order to be efficiently recruited to viral RNA.

**Figure 6:**
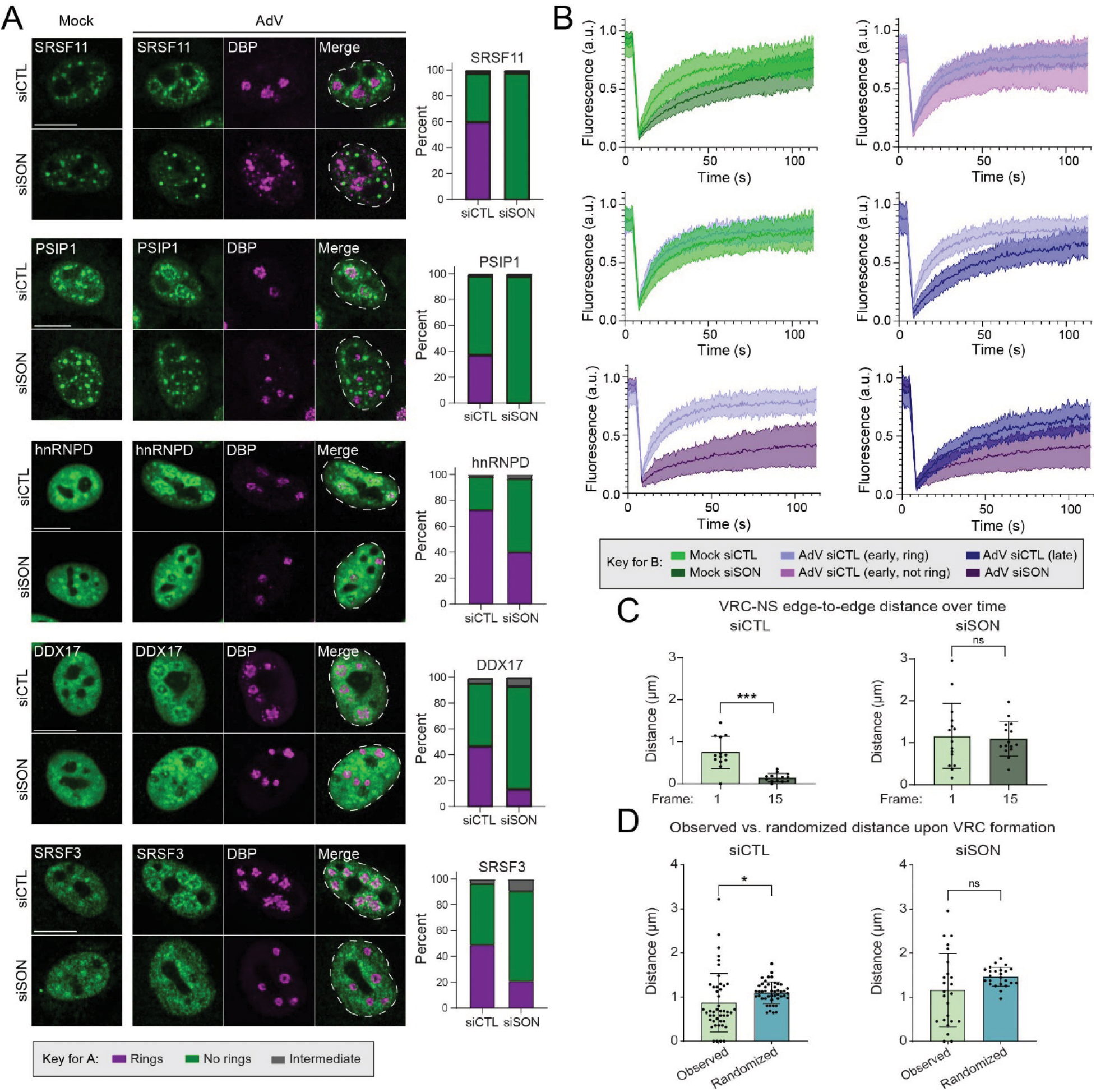
SON is required for efficient dynamic exchange of NS proteins and impacts VRC formation during infection. A) A549 cells were infected with AdV for 20 h following SON knockdown, then immunostained for proteins identified in the siRNA screen (left). Quantification of cells in the early stage of infection which demonstrate the formation of rings around VRCs (right). At least 30 cells were quantified per condition for each protein. B) Fluorescence recovery curves of NS in HeLa mScarlet3-SRRM2 cells. *n* = 2 independent replicates pooled. A total of 20 NS were analyzed for each condition except AdV siSON (19 speckles). C) Average edge-to-edge distances of VRCs and NS at frames 1 and 15 of live-cell imaging for siCTL cells (left) and siSON cells (right). Frame 1 represents the first frame in which VRCs were detected by thresholding, and frame 15 represents 3.5 h after frame 1. *n* = 13 cells (siCTL) and *n* = 15 cells (siSON). D) Observed vs. randomized edge-to-edge distances in siCTL cells (left) and siSON cells (right). Each data point represents a VRC. *n* = 46 VRCs (siCTL) and *n* = 24 VRCs (siSON). C-D) Significance was determined by a two-tailed paired t-test, with *p≤0.05, **p≤0.01, ***p≤0.001, ****p≤0.0001, ns=not significant. Graphs represent mean +/- standard deviation.

We reasoned that reduced recruitment of speckle proteins to VRCs during SON knockdown could be explained by limited ability of these components to exchange between the speckle and wider nucleoplasm. SON has been shown to positively regulate fluidity of nuclear speckles^70^. We therefore performed fluorescence recovery after photobleaching (FRAP) to estimate the exchange rate of components between bleached areas and the surrounding phase. We knocked down SON in HeLa mScarlet-SRRM2 cells for 24 h, then infected cells with AdV. At 18 hpi, we photobleached speckles, and assessed their recovery over a period of 104.5 s (**Figures 6B, S6B-D**). In agreement with previous work^70^, we found that SON knockdown in uninfected cells resulted in significantly slower rates of SRRM2 recovery (*t*_1/2_ = 39.1 s) compared to cells treated with nontargeting control siRNA (*t*_1/2_ = 15.4 s). We next analyzed speckles in infected cells. To account for the possibility that properties of speckles within early-stage AdV-infected cells adjacent to VRCs may differ from those not recruited to VRCs within a given cell, we analyzed these separately. We additionally distinguished between cells at early-stage of infection (using ring formation as an indicator) and late-stage cells, where speckles are larger and mostly at the nuclear periphery. For early-stage infections, we found no significant difference in rates of recovery between the speckles in rings (*t*_1/2_ = 11.6 s) and outside of rings (*t*_1/2_ = 15.7 s) (**Figures 6B, S6B-D**). Additionally, there was no significant difference between uninfected and early-stage infected cells treated with siCTL. In late-stage infected cells, we observed slower (though not significant) rates of recovery (*t*_1/2_ = 26.2 s) when compared to speckles in early-stage infections within VRC rings. This finding suggests that AdV induces slower dynamic exchange of SRRM2 (and possibly other resident proteins) as infection progresses. During knockdown of SON, this rate became significantly slower (*t*_1/2_ = 37.8 s), with a total recovery of 46.4% (**Figures 6B, S6B-D**). This total recovery was significantly less than the other conditions, which were comparable and ranged from 78.7% (mock siSON) to 85.8% (AdV siCTL early stage, not ring) (**Figure S6D**). These data demonstrate that SON is required for efficient dynamic exchange of SRRM2 with the wider nucleoplasm during infection.

We next investigated the dynamics of speckles and early-stage VRCs by performing live-cell imaging. We knocked down SON in HeLa mScarlet-SRRM2 cells for 24 h prior to infection with the previously reported AdGDBP (GDBP) virus^39^, which expresses DBP fused to EGFP to enable visualization of replication centers. We acquired images every 15 min starting at 8 hpi (**Supplemental movies 1-2**). To determine whether SON is required for efficient recruitment of speckles to VRCs, or vice versa, we analyzed the proximity of speckles and VRCs for 3.5 h immediately following VRC formation (**Figures 6C, S6E**). We found that as infection progressed, the distances between the edges of these two nuclear structures significantly decreased in the siCTL cells, but not in the siSON cells.

We then asked whether SON impacts the proximity relative to NS in which nascent VRCs initially appear. We measured the observed VRC-NS distances within the first frame where VRCs were detectable on a cell-by-cell basis, and calculated the minimum edge-to-edge distance of each VRC to the closest NS. We then performed 1000 rounds of randomization analysis, where all VRCs were randomly repositioned within the remaining nuclear area while the NS masks remained fixed, before measuring the randomized VRC-NS distances. This allowed us to determine whether VRCs initially form closer to speckles than would be expected by chance in the siCTL control cells when compared to the siSON cells. We found that the observed VRC-NS distances were indeed significantly reduced compared to the randomized distances in the siCTL cells, but not in the siSON cells, on both a VRC-level and cell-level basis (**Figures 6D, S6F**). These data collectively indicate that SON impacts both the initial formation of VRCs, as well as the proximity of VRCs and NS over the time course of infection. We propose a model where SON mediates viral RNA expression by facilitating the recruitment of NS to nascent VRCs as well as the dynamic exchange of NS proteins with the nucleoplasm (**Figure 7**).

**Figure 7:**
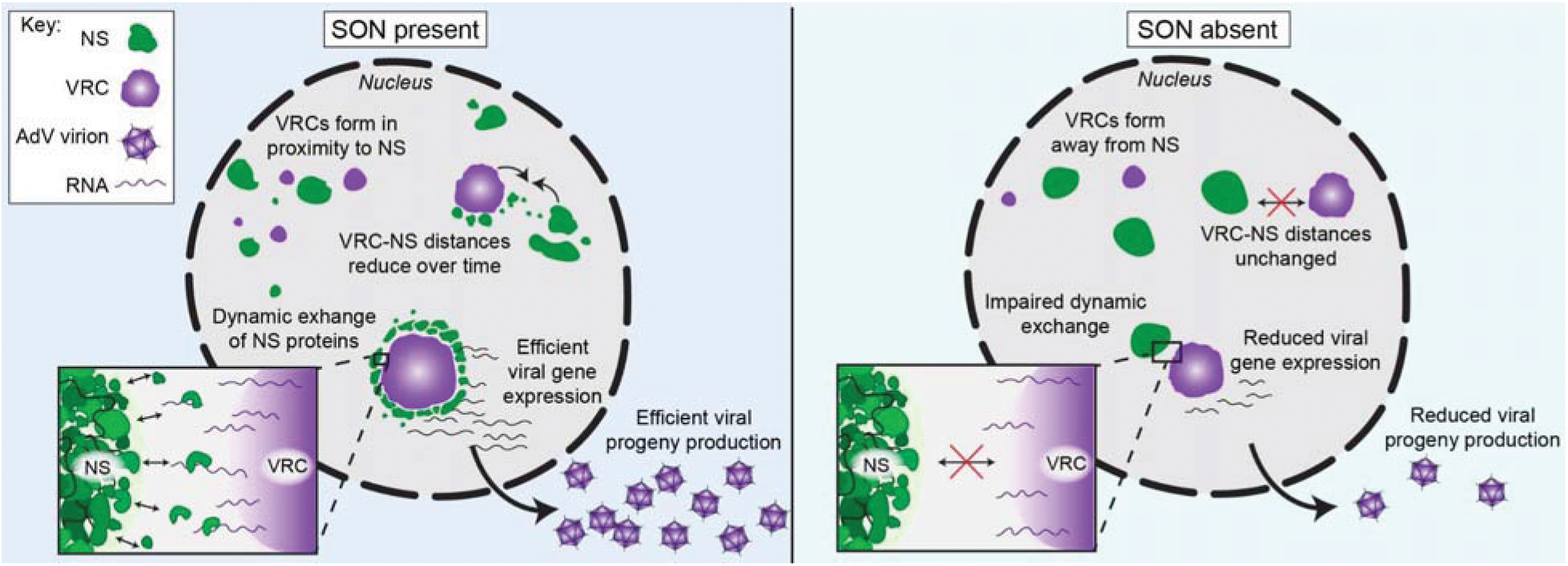
Model of the impact of SON on AdV infection. **Left panel:** When SON is present, VRCs form closer to NS than would be expected by chance, and the proximity of the two structures decreases over time. NS proteins exchange between the speckle and the wider nucleoplasm, leading to efficient viral gene expression and productive infection. **Right panel:** In the absence of SON, NS form away from VRCs and the distances between the structures remain unchanged. Certain speckle proteins are unable to freely exchange between the speckle and the nucleoplasm, resulting in reduced viral gene expression and impaired infection.

## DISCUSSION

AdV has long served as a powerful model system for investigating RNA biology. While it is known that this virus depends on cellular RNA-processing machinery, the scope of this dependency has been largely limited to candidate-based studies. Here we employed multiple unbiased screening approaches to provide a more comprehensive picture of how AdV reorganizes nuclear RBPs and co-opts proteins to promote infection. By combining optical pooled screening of a SPOTLITES library together with a targeted functional screen, we identified RBPs relocalized by AdV that are required for infection. This analysis revealed a requirement for multiple NS proteins and demonstrated that the core protein SON is a critical factor for promoting viral gene expression. This study demonstrates the utility of optical pooled screening for investigating virus-host interactions at the proteome-level, and specifically highlights a requirement of SON for viral infection.

Optical pooled screening is not yet widely used to study viral infections. While previous work has employed this approach to determine the impact of genetic perturbations on infection^23,24^, our study is the first to utilize this method to investigate the impact of viral infection on host protein dynamics at scale. Previous work has employed optical pooled screening of this type of cell library to investigate the unfolded protein response^20^. Here we expand the utility of this approach to include investigating viral infections, and we expect this approach can be readily applied to investigate other pathogens that remodel host cell architecture. Additionally, our results have implications for the investigation of diseases in which NS morphology is altered. A recent study found a strong correlation between altered NS localization patterns and patient outcomes for clear cell renal cell carcinoma^71^. Multiple additional studies have linked speckles with cancers and other disease states^72–75^, and the term “nuclear speckleopathies” has been recently coined to denote genetic disorders caused by mutations in genes encoding NS proteins^73^. We propose that our approach is well suited for studying changes in NS morphology, and for identifying factors which localize to these nuclear bodies in particular disease states. This method should provide a broad, powerful approach for investigating any other subcellular structure where protein composition is known to change in response to a given stressor.

Relocalization alone does not imply functional importance. By pairing the results from the optical pooled screen with those of a targeted siRNA screen, we were able to determine which proteins were both reorganized by AdV and had functional relevance to infection. These include factors involved in multiple aspects of RNA processing, including transcriptional regulation (SUPT5H, SUPT6H, PSIP1), 3’ end cleavage and polyadenylation (CSTF2), splicing (TRA2B, RBM22, multiple hnRNP and SRSF proteins), exporters (XPO1), and ribosome biogenesis (RBM28, NCL, UTP14A). Some of these proteins such as NCL and XPO1 have been previously shown to be involved with AdV infection^76–78^, but their roles remain incompletely understood. Additionally, AdV transcripts have long been used to study functions of hnRNP and SRSF splicing factors, and this virus is known to depend on proteins from these families to promote viral RNA processing^79–85^. However, the roles of many of the other identified proteins during AdV infection remain uncharacterized. Our findings exemplify the myriad cellular pathways co-opted by AdV, and raise further questions regarding how each protein is redirected to facilitate processing of viral RNA.

The finding that multiple NS proteins are relocalized by AdV and are also required for efficient progeny production is intriguing. While it has been known for decades that splicing factors form rings around early VRCs before transitioning to enlarged speckles at the nuclear periphery^25,35,36^, these studies primarily utilized antibodies recognizing common snRNP antigens and did not explore the scope of proteins that comprise this phenotype. Our findings build on these observations, demonstrating that these enlarged speckles include the two core proteins as well as numerous other RBPs. Additionally, we show that recruitment of splicing factors to rings around VRCs is dependent on the presence of SON. Interestingly, SON depletion results in reduced expression of many viral genes, but E1A expression was largely unaffected or enhanced. As the first viral gene expressed, E1A expression is initiated prior to extensive nuclear reorganization and formation of VRCs, so these results may reflect differential regulatory mechanisms of viral gene expression over the course of infection. We further expand on previous findings that snRNPs no longer form a ring pattern following transcription inhibition^86^ by demonstrating that DRB treatment prevents core protein SRRM2 from localizing to rings. Collectively our results establish a viral requirement for SON, and define a comprehensive list of RBPs recruited to rings adjacent to viral centers.

Our findings confirm non-redundant roles of NS core proteins. Previous work has shown that SON and SRRM2 organize distinct molecular networks within speckles, and that the multiphase formation and client partitioning of these nuclear bodies likely involves fundamentally distinct biochemical properties of the two proteins^42^. Prior FRAP experiments have shown that SRRM2 recovers relatively quickly following photobleaching, while SON recovers much more slowly, suggesting that SON imparts viscoelastic properties to speckles^87^. Here we found that depletion of SRRM2 had negligible impact on any aspect of the viral replication cycle and did not affect the ability of SON to form rings around VRCs. In contrast, SON depletion impacted viral gene expression, prevented SRRM2 from localizing to rings, and altered the proximity of speckles and VRCs over time. We also found that SON depletion resulted in slower fluorescence recovery of SRRM2 during infection, but whether this occurrence is applicable to other RBPs requires further investigation. NS are dynamic structures, and their components are in continuous flux with the nucleoplasm^88,89^. SON itself is essential for NS organization^67^. It is possible that disrupting this organization alters internal dynamics of speckles in such a way that resident proteins can no longer exchange with the nucleoplasm as effectively. Furthermore, a recent study identified an interchromatin network of connections that functionally links NS throughout the nucleus and coordinates speckle protein trafficking^90^. We find it likely that the depletion of SON disrupts this trafficking, thereby precluding NS proteins from accessing nascent VRCs and viral RNA. Viral systems such as AdV present promising tools for investigating speckle properties and functions.

Several unanswered questions remain regarding the role(s) of NS during AdV infection. First, while our work demonstrates a requirement of AdV for a NS core protein, the question of whether the virus co-opts these nuclear bodies and/or their components by a particular strategy remains unanswered. Clues from the literature suggest that viral manipulation of speckle protein phosphorylation may be one such strategy. Specifically, speckle protein SRSF1 is known to be dephosphorylated by protein phosphatase 2A (PP2A) following redirection by AdV protein E4orf4^85^, and redistribution of this protein to enlarged speckles has been shown to correlate with splicing of certain viral transcripts^91^. Furthermore, SR proteins purified from late AdV-infected cells have been shown to be inactivated as splicing enhancer or splicing repressor proteins by virus-induced dephosphorylation^92^. Since regulation of SR protein phosphorylation is known to mediate localization to and from speckles^93–100^, it is tempting to speculate that redirection of kinases and/or phosphatases may be one strategy by which AdV facilitates expression and processing of viral transcripts by temporally manipulating speckle component localization. Second, our data demonstrate a distinction between the fluorescence recovery rate of speckles during different stages of infection, but whether this finding reflects an infection stage-specific change in function for speckles remains unknown. Previous work has identified that spliced exons of AdV late transcripts colocalize with enlarged speckles in the late stage, which led to the proposal that these structures could be involved with RNA export^59^, but this needs to be experimentally tested. Additionally, it remains unclear the extent to which the virus mediates formation of these larger structures, or if their formation is a passive consequence of rearrangements to nuclear architecture during the late stage. Lastly, we found several infection-relevant proteins which cannot relocalize to rings in the absence of SON, but we do not yet know the scope of this impaired relocalization. Future work investigating common features of proteins which relocalize in the presence or absence of SON, may reveal clues to help explain how SON mediates NS organization and protein mobility.

## Limitations

It is possible that the process of tagging affects normal localization patterns of proteins in the SPOTLITES library, which may have affected the results. Additionally, not every gene tolerated insertion of the tag, meaning that the final cell library may have underrepresented the known repertoire of RBPs. It is therefore likely that additional proteins may relocalize upon infection but were not identified in this study. AdV infection also alters the host nuclear environment, so it is possible that this impacted the ability of the deep learning-based analysis to recognize common phenotypes (i.e. nucleoli) between uninfected and infected cells, even if subcellular localization of a given protein was unchanged. We recognize that SON is an essential protein and its knockdown can induce cell cycle arrest. While our imaging and FRAP analyses of speckles were performed in cells exhibiting normal morphology, future studies should consider alternative strategies to minimize detrimental effects on the cell. Lastly, while double knockdown of SON and SRRM2 has been previously utilized as an approach to perturb speckles^60,66,87^, we cannot draw any conclusions regarding the impact of the entire nuclear body on infection given the striking impact of SON alone.

## ACKNOWLEDGEMENTS

We thank members of the Weitzman, Shalem, and Price Labs for insightful discussions and input, and for their thoughtful feedback and careful reading of the manuscript. We are grateful to A. Mehle, A. Levine, and P. Hearing for generous gifts of reagents. We thank the UPenn Cell and Developmental Biology Microscopy Core and The Wistar Institute’s Imaging Shared Resource for imaging assistance and technical support. Funding support for The Wistar Institute core facilities was provided by Cancer Center Support Grant P30 CA010815. This research was supported by NIAID grants R01-AI145266 (M.D.W.), R01-AI89522 (M.D.W. and O.S.), and R00-AI159049 (A.M.P.). M.D.W. was supported in part by the Arthur Vincent Meigs Endowed Chair in Pediatrics. Additional support came from the National Institutes of Health (R35GM142837, D.C.), American Cancer Society Research Scholar Grant (RSG-25-1432735-01-CCB to D.C), Johns Hopkins Catalyst Award (D.C.), Individual National Research Service Award F31-AI181460 (A.R.N.A.), and a CIHR post-doctoral fellowship (T.M.T.). The funders had no role in study design, data collection and analysis, decision to publish, or preparation of the manuscript.

## AUTHOR CONTRIBUTIONS

A.R.N.A., T.L., O.S., A.M.P., and M.D.W. conceived the study, designed the experiments, and interpreted experimental data. T.M.T. provided additional input into study design. A.R.N.A., T.L., H.C., and E.H. performed experiments. T.L. analyzed the high-throughput microscopy and *in situ* sequencing experiment and performed the deep learning analysis. S.Y. performed analysis of live-cell imaging. G.P. assisted with immunofluorescence analysis. D.C., O.S., A.M.P., and M.D.W. supervised the work and acquired funding. A.R.N.A., T.L., O.S., A.M.P., and M.D.W. wrote the manuscript. All authors read, edited, and approved the final paper.

## METHODS

### Cell culture

All cell lines were obtained from and authenticated by the American Type Culture Collection (ATCC). Cells were cultured under standard conditions (37°C and 5% CO_2_) in sterile, TC-treated, nonpyrogenic, polystyrene tissue culture dishes (Corning, 430167). SPOTLITE library cells were generated as previously described^20^. HeLa mScarlet3-SRRM2 cells were a gift from D. Cai. A549 (ATCC, CCL-185), HEK293 (ATCC, CRL-1573), and HeLa (ATCC, CCL-2) cells were cultured in DMEM (Corning, 10-013-CV) supplemented with 10% FBS (VWR, 89510-186) and 1% Pen/Strep (Gibco, 15140-122). All cell lines tested negative for mycoplasma using the LookOut Mycoplasma PCR Detection Kit (Sigma-Aldrich, MP0035-1KT). Mycoplasma testing was carried out twice a year.

### DRB treatment

5,6-Dichlorobenzimidazole riboside (DRB) (MedChem Express, HY-14392-10mg), was added at 60 □M for 2 h following 18 h of infection.

### Viruses and Infections

WT Ad5 (ATCC, VR-5) was propagated on HEK293 cells, purified using two sequential rounds of ultracentrifugation in cesium chloride density gradients, and stored in dilution solution (8 mM Tris pH 8, 80 mM NaCl, 0.08% BSA, and 40% glycerol) at −20 °C (short term) or −80 °C (long term). Viral titer was determined by plaque assay on HEK293 cells. All infections with WT Ad5 were performed at a multiplicity of infection (MOI) of 10 and harvested at indicated hours post infection (hpi). Infections were performed by diluting virus in serum-free growth media and incubating cells for 2 h at 37°C with gentle rocking every 20 min. After 2 h, viral inoculum was replaced with full-serum growth media. AdNTE and AdGDBP were gifts from A. Mehle. For infections with AdNTE, cells were inoculated with virus diluted in DMEM without FBS at MOI 5 as previously described^39^. The inoculum was removed after 1 h and replaced with DMEM containing 10% FBS.

### High throughput microscopy phenotyping and in situ sequencing

SPOTLITE library cells were seeded on glass-bottom 6 well plates (CellVis, P06-1.5H-N) at 40% confluency, then infected the next day with WT Ad5 (see “Viruses and Infections”). At 23.5 hpi, medium was aspirated and replaced with fresh medium containing JFX549-HaloTag ligand (Promega, GA1110) at 50 nM. Cells were incubated for 15 min, washed 3 X for 5 min with fresh medium, then fixed at 24 hpi. Cells were then processed as previously published^20^. Cells were imaged on an inverted Ti2 Nikon Eclipse microscope. Phenotyping was performed by imaging DAPI, G3BP1, and Halotag channels at 40X magnification. The incorporation, imaging, and cleavage steps were repeated for a total of 14 cycles. Every genotyping cycle was performed by imaging DAPI and each of the 4 nucleotide channels at 10X magnification.

### Training the variational autoencoder Cytoself

The variational autoencoder (VAE) was trained on 320,184 cropped single-cell images derived from Cellpose-based segmentation of the G3BP1 channel, using either the HaloTag fluorescence channel or DBP staining as model input. To enable robust learning of subcellular localization phenotypes, a semi-manually curated training dataset was assembled containing 13 distinct localization classes, including both canonical cellular compartments and VRCs. The training dataset for VRCs was generated by infecting HeLa cells expressing G3BP1 endogenously tagged with mClover3 with WT Ad5 (see “Viruses and Infections”) for 16 or 24 h before fixing and immunostaining for DBP. These timepoints were chosen to capture a range of VRC phenotypes. Images were captured on an inverted Ti2 Nikon Eclipse microscope using a motorized programmable stage. Following segmentation, approximately 10,000 - 30,000 cells were selected to generate the training dataset for each of the 13 distinct localization patterns. By incorporating examples of normal cellular organization alongside infection-associated structures, the VAE was trained to encode diverse localization phenotypes within a shared 64-dimensional latent space. The model learned compressed latent representations that captured spatial localization features across individual cells while preserving phenotypic relationships between classes, enabling discrimination of both subtle and pronounced localization differences associated with viral infection.

### siRNA screen

The siRNA library was designed using the Horizon Discovery Cherry-Pick Custom Library Tool, using ON-TARGETplus SMARTpool reagents so that each gene of interest was targeted by four pooled siRNAs. A total of 80 RBPs were chosen for the screen based on silhouette score, and at least one RBP from each group of common relocalization patterns (**Figure 2B**) was selected. POLR2A was additionally included as a positive control. A549 cells were plated at 10,000 cells/well onto flat-bottom 96-well plates (Genesee Scientific, 25-109), and PBS was added to the outside wells to minimize evaporation over the course of the experiment. Cells were transfected the next day with the siRNA library for a final concentration of 1 pmol/well. siRNAs were diluted in Opti-MEM (Gibco, 31985088), then added to an equal volume of Opti-MEM containing Lipofectamine RNAiMAX Transfection Reagent (Thermo-Fisher, 13778150) following the manufacturer’s protocol. siRNA-lipid complexes were incubated for 5-15 minutes at room temperature, then added to the wells for a final volume of 0.3 uL RNAiMAX and 10 uL siRNA-lipid complex per well. After 24 h, cells were infected with AdNTE (see “Viruses and Infections”), then harvested at 24 and 48 hpi for analysis of viral progeny. At indicated timepoints, the plates underwent 3 cycles of freeze/thaws using liquid nitrogen and a 37°C water bath, then were spun at 500 rpm for 10 min to pellet cell debris. Supernatant was diluted in serum-free DMEM to reinfect new cells seeded in white 96-well flat-bottom plates (Corning, 3917). Following reinfection, nanoluciferase activity was measured at 6 hpi using NanoGLO reagents (Promega, N1110) following the manufacturer’s instructions. Luciferase activity was determined on a FLUOstar Omega microplate reader with software version 5.50 R3. The RLU values were blank-adjusted, then compared to the non-targeting siRNA control wells. Four wells were analyzed for each siRNA, and each plate contained an internal negative control to minimize batch effects. The screen was then repeated with each siRNA in a different location on the corresponding plate to ensure reproducibility. The Log_2_FC of the RLU values for each siRNA relative to the non-targeting control were calculated, and results were plotted using GraphPad Prism 10.2.2.

### Viability assay

A549 cells were transfected with the siRNA library in quadruplicate in white 96-well flat-bottom plates (Corning, 3917) for 72 h, then treated with CellTiter Glo 2.0 reagent (Promega, G9241) following manufacturer’s instructions to analyze ATP production as a metric for cell viability.

Each plate included an untransfected negative control, and cells treated with 1 □M bortezomib (MedChem Express, HY-10227) as a positive control. Luciferase activity was determined on a FLUOstar Omega microplate reader with software version 5.50 R3 as described above, and RLU values relative to the untransfected cells were determined.

### siRNA transfections

All siRNA transfections were performed using the standard protocol for Lipofectamine RNAiMAX (Thermo-Fisher, 13778150). The following siRNA pools were obtained from Dharmacon: SUPT5H (L-016234-00-0005), SUPT6H (L-010540-00-0005), SRSF3 (L-030081-00-0005), hnRNPK (L-011692-00-0005), SFPQ (L-006455-00-0005), SRRT (L-019234-01-0005), SON (L-012983-00-0005), SRRM2 (L-015368-00-0005), non-targeting control (D-001810-10-05). The complete list for the siRNA screen is provided as a supplementary file.

### Immunoblotting

Protein samples were prepared using lithium dodecyl sulfate (LDS) loading buffer (Thermo-Fisher, NP0007) supplemented with 1% 2-mercaptoethanol (BME) and boiled at 95°C for 10 min. Protein lysates were separated by SDS-PAGE and transferred onto methanol-activated polyvinylidene fluoride (PVDF) membranes (Millipore Sigma, IPFL00010). For analysis using SON or SRRM2 antibodies, transfers were performed overnight at 30V. For analysis using all other antibodies, transfers were performed for 90 min at 40V. Membranes were blocked for 1 h in blocking buffer (3% BSA in TBST for analysis of phosphorylated proteins, 5% milk in TBST supplemented with 0.05% sodium azide for all others), then incubated in primary antibodies diluted in blocking buffer overnight at 4°C. Membranes were washed 3 X for 5 min in TBST, incubated in HRP-conjugated secondary antibodies diluted in TBST for 1 h, then washed again 3 X for 5 min in TBST. Membranes were incubated for 2 min in Pierce SuperSignal West Pico PLUS Chemiluminescent Substrate (Thermo-Fisher, 34580) or SuperSignal West Femto Maximum Sensitivity Substrate (Thermo-Fisher, 34095) before protein detection using a Syngene G-Box with GeneSys acquisition software. Images were assembled in Adobe Illustrator 28.4.

### Antibodies

The following antibodies for viral proteins were used: Anti-adenovirus Type 5 antibody raised against whole adenovirus capsids (Abcam, ab6982, rabbit, polyclonal, WB 1:5000), DBP (gift from A. Levine, mouse, clone B6-8, WB 1:1000, IF 1:400), DBP (Cusabio, CSB-PA365892ZA01HIL, rabbit, polyclonal, IF 1:500), E1A (BD Biosciences, 554155, mouse, clone M58, WB 1:1000), E4orf6 (Gift from P. Hearing, mouse, clone RSA3, WB 1:500).

The following antibodies for cellular proteins were used: GAPDH (GeneTex, GTX100118, rabbit, polyclonal, WB 1:5000), SON (Abcam, ab317311, rabbit, clone EPR28617-69, WB 1:1000, IF 1:500), SRRM2 (Abcam, ab317817, rabbit, clone EPR28620-54, WB 1:1000, IF 1:500), hnRNPD (Abcam, ab61193, rabbit, polyclonal, IF 1:500), hnRNPU (Abcam, ab20666, rabbit, polyclonal, IF 1:500), PNN (Novus Biologicals, NBP1-87784, rabbit, polyclonal, IF 1:500), FUBP1 (Abcam, ab181111, rabbit, clone EPR12327, IF 1:500), LUC7L2 (ThermoFisher, A304-540A-T, rabbit, polyclonal, IF 1:500), SRSF3 (ThermoFisher, 33-9400, mouse, clone 1H4G7, IF 1:500), SRSF3 (ThermoFisher, 33-420-0, mouse, clone 7B4 (7B4A12), WB 1:1000), Pol II p-Ser2 (Abcam, ab5095, rabbit, polyclonal, IF 1:500), SRSF11 (Antibodies Online, ABIN486013, rabbit, polyclonal, IF 1:500), PSIP1 (Abcam, ab177159, rabbit, clone EPR11890, IF 1:500), DDX17 (Santa Cruz, sc-27112, mouse, clone C-9, IF 1:500), SUPT5H (Invitrogen, PA5-56100, rabbit, polyclonal, WB 1:1000), SUPT6H (Proteintech, 23073-1-AP, rabbit, polyclonal, WB 1:1000), hnRNPK (Abcam, ab39975, mouse, clone 3C2, WB 1:1000), SFPQ (Abcam, ab38148, rabbit, polyclonal, WB 1:1000), SRRT (Bethyl Laboratories, A304-551A-T, rabbit, polyclonal, WB 1:1000).

For immunoblot analysis, horseradish peroxidase-conjugated (HRP) goat anti-mouse (Jackson Laboratories, 115-035-003) or goat anti-rabbit (Jackson Laboratories, 111-035-045) secondary antibodies were used at a concentration of 1:5000. For immunofluorescence, Alexa Fluor fluorophore-conjugated secondaries were used at a concentration of 1:1000. Fluorophores used were goat anti-rabbit 488 (Thermo-Fisher, A-11008), goat anti-mouse 488 (Thermo-Fisher, A-11001), goat anti-rabbit 647 (Thermo-Fisher, A-21245), goat anti-rabbit 568 (Thermo-Fisher, A-11011), and goat anti-mouse 568 (Thermo-Fisher, A-11004).

### Immunofluorescence and image analysis

For microscopy experiments other than the optical pooled screen, cells were grown on 12-mm glass coverslips (Electron Microscopy Sciences, 72196-12) in 24-well plates. Cells were transfected and/or infected as indicated, then fixed with 4% paraformaldehyde (PFA) in PBS for 15 min before 2 X washes in PBS. Cells were permeabilized with 0.5% Triton-X in PBS for 10 min, washed 2X in PBS, then blocked in 3% bovine serum albumin (BSA) in PBS (+ 0.05% sodium azide) for 1 h. Samples were then incubated with primary antibodies in 3% BSA in PBS (+ 0.05% sodium azide) for 1 h, washed 3X in 3% BSA in PBS, incubated with secondary antibodies and 4,6-diamidino-2-phenylindole (DAPI) for 1 h, then washed 3X in PBS. Coverslips were mounted onto glass slides with ProLong Gold Antifade Reagent (Cell Signaling Technologies, 9071) and immunofluorescence was visualized on a Zeiss LSM 980 Confocal microscope (Cell and Developmental Microscopy Core at Penn) with ZEN v2.3 acquisition software (all confocal images), or Leica DMi8 microscope with LAS X Software version 3.9.1.28433 (for verification of infection efficiency for the optical pooled screen). Images were acquired for at least 3 fields of view per coverslip. Images were processed using FIJI (v1.54p) and Adobe Illustrator (v28.4.1). For quantification of ring formation, a minimum of 30 cells per condition were assessed by blinded analysis.

### Fluorescence recovery after photobleaching

HeLa mScarlet3-SRRM2 cells were infected with WT AdV for 18 h following the indicated knockdown. Live cell imaging for FRAP was performed on a Leica Stellaris 3X TauSTED microscope with LAS-X software and the 63X/NA 1.4 objective. All imaging of samples was conducted under standard conditions of 37°C and 5% CO_2_. Fluorescence was monitored for 5 s prior to photobleaching, and recovery was monitored for 104.5 s after photobleaching. Fluorescence recovery was analyzed in GraphPad Prism 10.2.2 using nonlinear regression and the one phase association model. Two independent replicates were pooled together for a total of 20 NS analyzed for each condition, except for AdV siSON (19 speckles).

### Live-cell microscopy and image processing

#### Infection and image acquisition

HeLa mScarlet3-SRRM2 cells cultured in 35 mm glass bottom dishes (MatTek, P35G-1.5-14-C) were transfected with siCTL or siSON for 24 h, then infected with GDBP at 10 MOI for 8 h. Media was replaced with FluoroBrite DMEM (Gibco, A18967-01) supplemented with 10% FBS and 1% Penn/Strep. NucBlue Live Cell Stain ReadyProbes (invitrogen, R37605) was added to visualize nuclei. Cells were imaged every 15 min on a Nikon Ti automated inverted microscope using a 40X objective and NIS-Elements software for 16.5 h; however, challenges due to photobleaching of mScarlet precluded analysis past ∼4 h. All imaging was conducted under standard conditions of 37°C and 5% CO_2_.

#### Nearest edge-to-edge distance calculation between VRC and NS

Time-lapse images of GDBP-infected cells were analyzed to determine the spatial relationship between VRCs and NS using custom python scripts. VRCs were segmented using Otsu thresholding, with an additional minimum pixel-intensity cutoff of 1000 to exclude low-intensity background signal. NS were segmented using three-class Multi-Otsu thresholding to distinguish bright NS signal from diffuse nucleoplasmic signal and dark background. To prevent unreliable segmentation at later time points due to photobleaching, NS masking was discontinued when the calculated Multi-Otsu threshold fell below 120. Otsu thresholding was also applied to the nuclear channel, followed by hole filling to generate continuous nuclear masks.

Edge-to-edge distances between segmented VRCs and NSs were subsequently quantified at each time point. For each source VRC, the minimum edge-to-edge distance from VRC to the nearest NS was calculated using a Euclidean distance transform. If the VRC and NS masks overlapped, the distance was defined as 0. Distances were then converted from pixels to micrometers using a calibration of 6.2066 pixels per micrometer. For each VRC, the minimum edge-to-edge distance to every NS in the same cell was calculated, and the smallest edge-to-edge distances were recorded. Finally, data from the first time point at which VRCs became detectable and the subsequent 3.5 hours of time points was pooled for statistical analysis.

#### Randomization analysis

To determine whether the observed VRC-NS proximity differed from that expected by chance, we also calculated randomized VRC-NS distances by spatially repositioning VRCs while keeping the nuclear and NS masks fixed. For each cell, the image corresponding to the first time point at which VRCs became detectable was used as the basis for randomization and comparison with the observed distances. The permitted randomization area was defined as the nuclear area excluding NS-occupied areas. Each VRC was repositioned while preserving its shape and orientation. When multiple VRCs were present in the same cell, all VRCs were repositioned in each round of randomization. A total of 1000 rounds of randomizations were performed for each cell.

For each round of randomization, the minimum distances between randomized VRCs and their closest NS were calculated using the same method described above. Therefore, each randomized VRC was associated with a distribution of 1,000 randomized VRC-NS distances, from which the median number was used for comparison with the corresponding observed VRC-NS distance. Analyses were performed both at per-VRC level and at per-cell level. For per-VRC analyses, each VRC represented an individual data point. For per-cell analyses, distances from all VRCs detected within the same cell were averaged to generate a single value per cell.

### Viral DNA replication qPCR

For analysis of viral genome replication, DNA was harvested from A549 cells following transfection and/or infection at the indicated timepoints. Cells were harvested by scraping, pelleted by centrifugation, and flash frozen. DNA extraction was performed using the PureLINK Genomic DNA Mini Kit (Invitrogen, Cat# K182002) using the manufacturer’s instructions. qPCR was performed with primers for the DBP genomic region and normalized to cellular tubulin. Samples were further normalized to the 4 hpi timepoint of siControl using the threshold cycle (ΔΔ*C_T_*) method via SYBR green PCR master mix (Thermo-Fisher, 4367659). qPCR was performed on a QuantStudio 7 Flex real-time PCR system (Applied Biosystems) using the manufacturer’s instructions for standard speed reactions. The primers used were as follows:

DBP forward: GCCATTGCGCCCAAGAAGAA
DBP reverse: CTGTCCACGATTACCTCTGGTGAT
Tubulin forward: CCAGATGCCAAGTGACAAGAC
Tubulin reverse: GAGTGAGTGACAAGAGAAGCC

### RNA isolation and RT-qPCR

Total RNA was isolated from cells by RNeasy Mini kit (Qiagen, 74106), following the manufacturer’s instructions. RNA was treated on-column with RNase-free DNase I (Qiagen, 79254), before converting to complementary DNA (cDNA) using 1 μg of input RNA in the High Capacity RNA-to-cDNA kit (Thermo-Fisher, 4387406). qPCR was performed using the standard protocol for SYBR Green reagents (Thermo-Fisher, 4367659) in a QuantStudio 7 Flex Real-Time PCR System (Applied Biosystems). All values were normalized by the threshold cycle (ΔΔ*C_T_*) method by normalizing first to internal controls. The primers used were as follows:

E1A (spliced) forward: ACGGTTGCAGGTCTTGTCATT
E1A (spliced) reverse: GTTCAGACACAGGACTGTAGACA
E1A (unspliced) forward: ACGGTTGCAGGTCTTGTCATT
E1A (unspliced) reverse: ACTCTATCACCCACTGCCCAT
DBP (forward): GCCATTGCGCCCAAGAAGAA
DBP (reverse): CTGTCCACGATTACCTCTGGTGAT
E4 forward: GACAGGAAACCGTGTGGAATA
E4 reverse: CACAGAGTACACAGTCCTTTCTC
TPL (spliced) forward: TTCCGCATCGCTGTCTG
TPL (spliced) reverse: CCGATCCAAGAGTACTGGAAAG
TPL (unspliced) forward: GTCCAGGGTTTCCTTGATGAT
TPL (unspliced) reverse: CCGATCCAAGAGTACTGGAAAG
Fiber forward: GAAAGGCGTCTAACCAGTCA
Fiber reverse: AAAGGCACAGTTGGAGGAC
ActB forward: CATGTACGTTGCTATCCAGGC
ActB reverse: CTCCTTAATGTCACGCACGAT

### Metabolic labeling of RNA

Nascent transcription was analyzed by labeling RNA with 200 M 4sU for 1 h in A549 cells following knockdown of SON. RNA was harvested using Trizol (ThermoFisher, 15596026) at 24 hpi following the manufacturer’s instructions. A fraction of the total RNA was reserved, then the remaining 4sU-labeled RNA was biotinylated with MTSEA-Biotin-XX (Biotium, 90066) as previously described^101^. Nascent RNA was separated from unlabeled using MyOne C1 Streptavidin Dynabeads (Thermo Fisher Scientific, 65-001). Biotin was removed using 100 mM dithiothreitol (DTT), and RNA was isopropanol precipitated. One µg of total RNA and an equivalent volume of nascent RNA were converted to cDNA. qPCR was performed and the relative transcription rates determined by the ΔΔCt method normalized to nascent Actin RNA.

### Plaque Assay

Cells were transfected and/or infected as indicated and harvested by scraping and pelleting at the indicated times post-infection. Samples were frozen and thawed 3 X using liquid nitrogen and a 37°C water bath. Centrifugation at maximum speed (21,130 × *g*) for 5 min at 4°C was performed to remove cell debris. Lysates were serially diluted in DMEM containing 1% Pen-Strep to infect confluent monolayers of HEK293 cells seeded in 12-well plates. At 2 hpi, infection medium was aspirated and cells were overlaid with DMEM containing 0.45% SeaPlaque agarose (Lonza, 50101), 2% FBS, and 1% Pen-Strep. At 5 days post-infection, 10% trichloroacetic acid was added to each well for 30 min at room temperature. The medium was aspirated, and plaques were stained with 1% crystal violet in 70% methanol for 10 min.

---

**Supplemental Figure 1:**
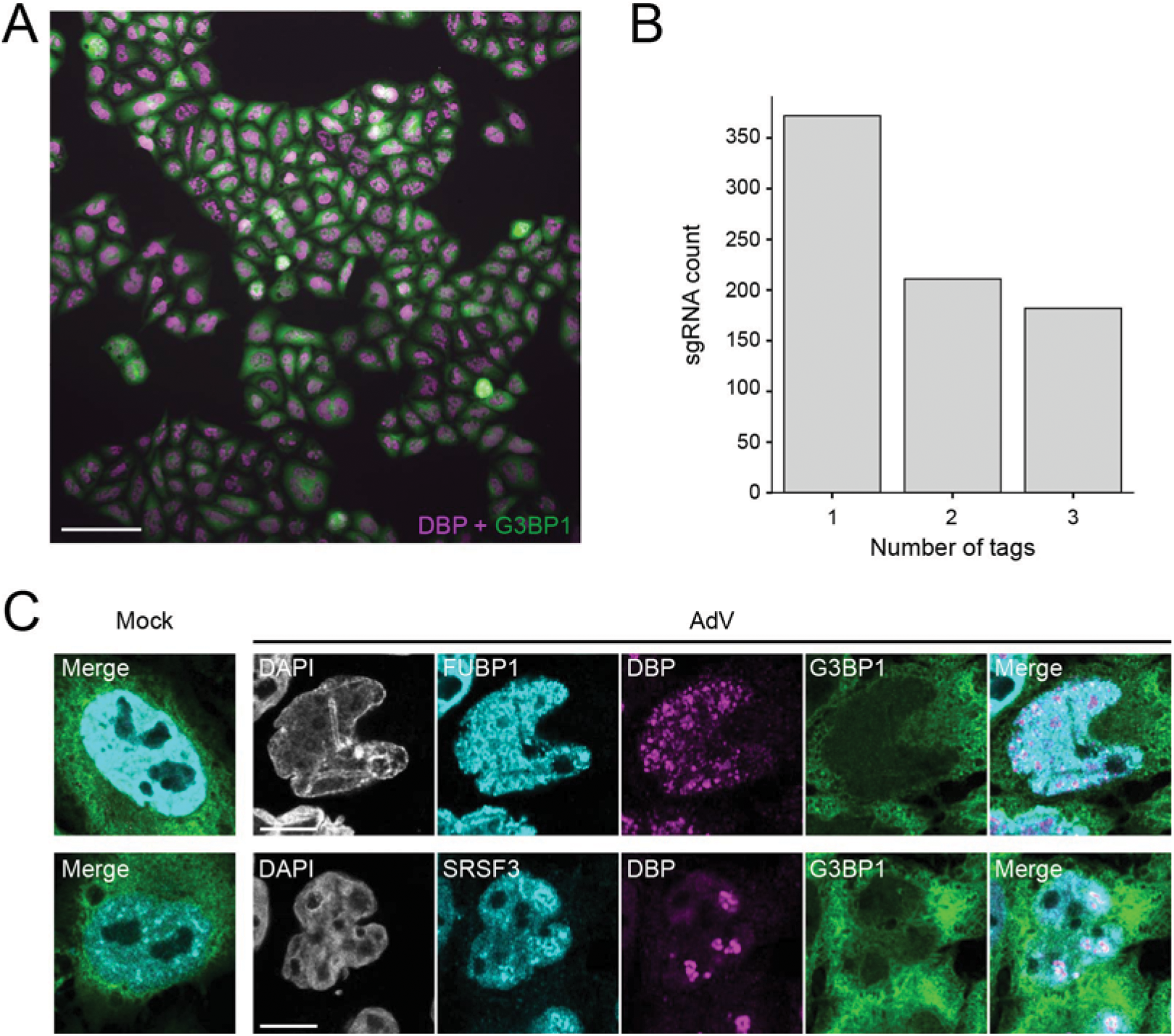
Validation of localization patterns identified in the optical pooled screen. A) Representative field of view of the SPOTLITES library cells infected in parallel to the optical pooled screen. DBP (magenta) is the viral DNA binding protein that marks sites of viral replication centers. G3BP1 (green) tagged with mClover3 acts as a cellular marker. Scale bar = 100 µm. B) Number of tags corresponding to each RBP. C) Confocal images of HeLa cells expressing endogenous G3BP1 tagged with mClover3 infected with AdV for 24 h, then immunostained for the indicated protein (cyan) to validate results of the optical pooled screen. Scale bar = 10 µm.

**Supplemental Figure 2:**
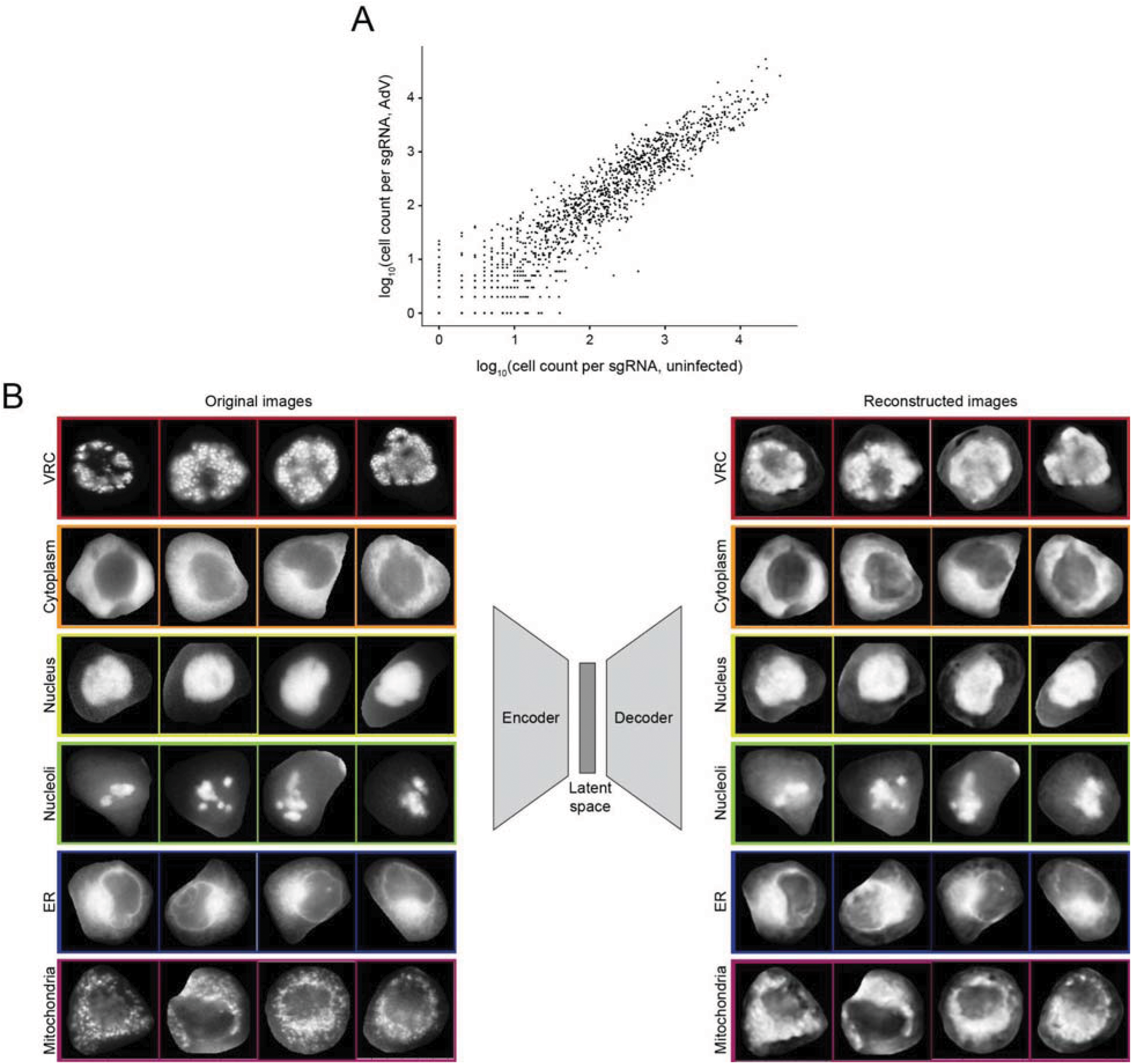
Training a deep learning algorithm to recognize VRCs. A) Comparison of sgRNA counts between uninfected and infected cell datasets. B) Original and reconstructed training images demonstrating viral (VRC) and example host (cytoplasm, nucleus, nucleoli, ER, mitochondria) localization patterns.

**Supplemental Figure 3:**
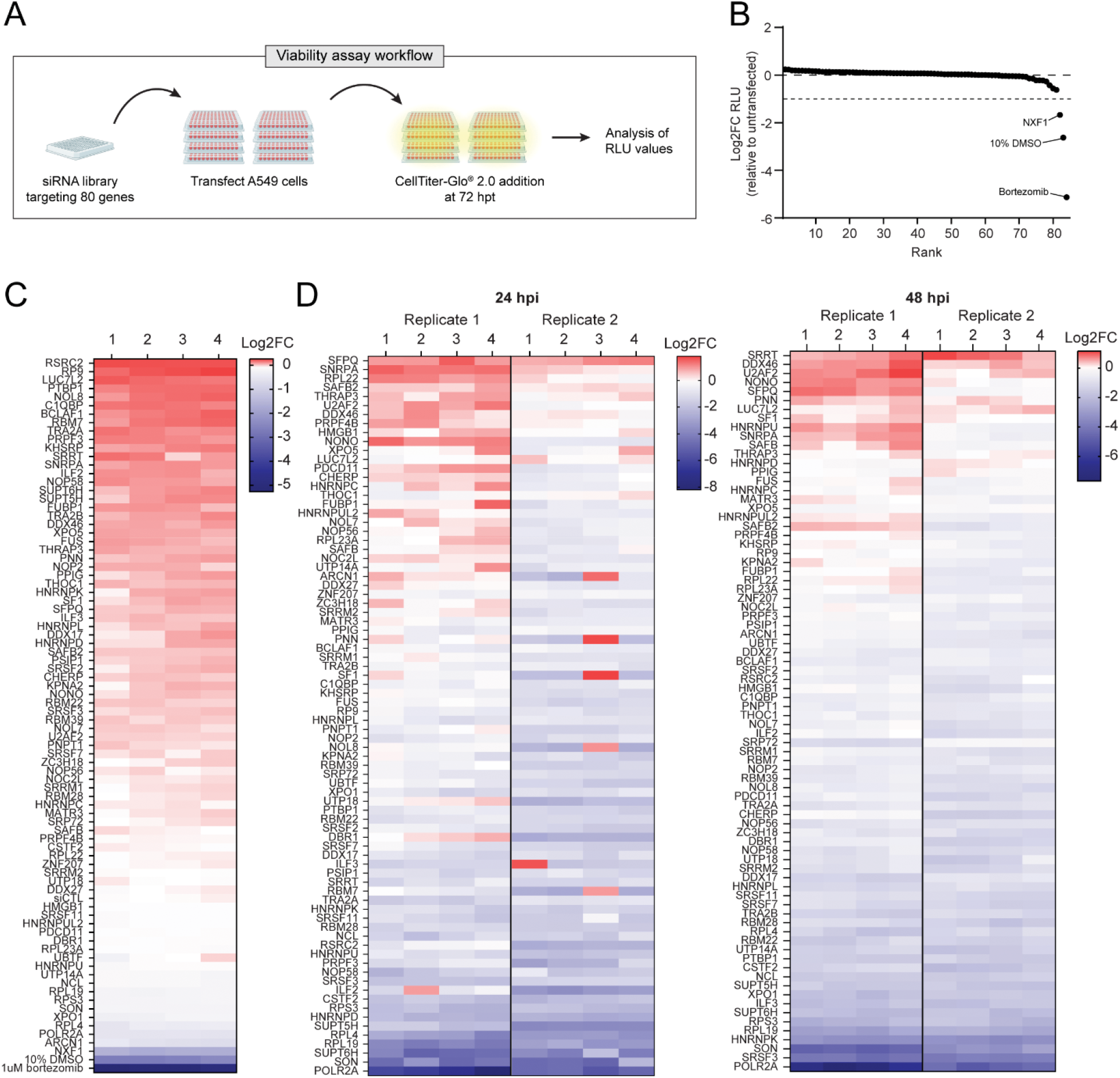
Analysis of cell viability following expression of siRNA library. A) Schematic for cell viability assay following expression of siRNA library. B) Treatments ranked by Log2FC blank-adjusted RLU values relative to untransfected cells. Treatment with 10% DMSO and 1uM bortezomib were positive controls to induce cell death. Only one siRNA (corresponding to NXF1) induced a Log2FC<-1, and so was excluded from analysis of the main siRNA screen. C) Heatmap of the Log2FC blank-adjusted RLU values of each well for the viability assay relative to untransfected cells. D) Heatmaps of the Log2FC blank-adjusted RLU values for each well relative to the nontargeting siCTL corresponding to the main siRNA screen. Values for both biological replicates of the screen are shown, corresponding to samples harvested at 24 hpi (left) and 48 hpi (right). Genes were ranked by the average Log2FC RLU across all 8 wells.

**Supplemental figure 4:**
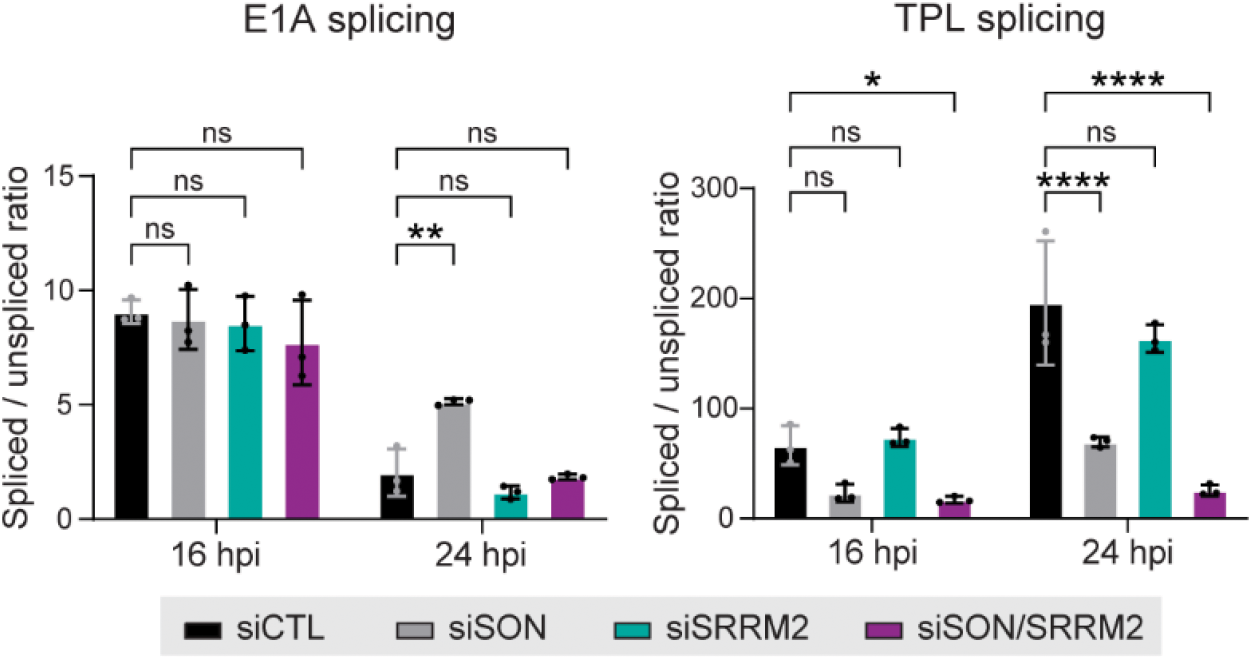
AdV splicing efficiency following knockdown of NS core proteins. Splicing efficiency determined by the relative ratio of spliced to unspliced transcripts of E1A and TPL. A549 cells were infected with AdV following depletion of SON and/or SRRM2, then transcripts analyzed by RT-qPCR. Significance was determined by two-way ANOVA with Dunnett’s multiple comparisons test. *p≤0.05, **p≤0.01, ***p≤0.001, ****p≤0.0001, ns=not significant. Graphs represent mean +/- standard deviation.

**Supplemental figure 5:**
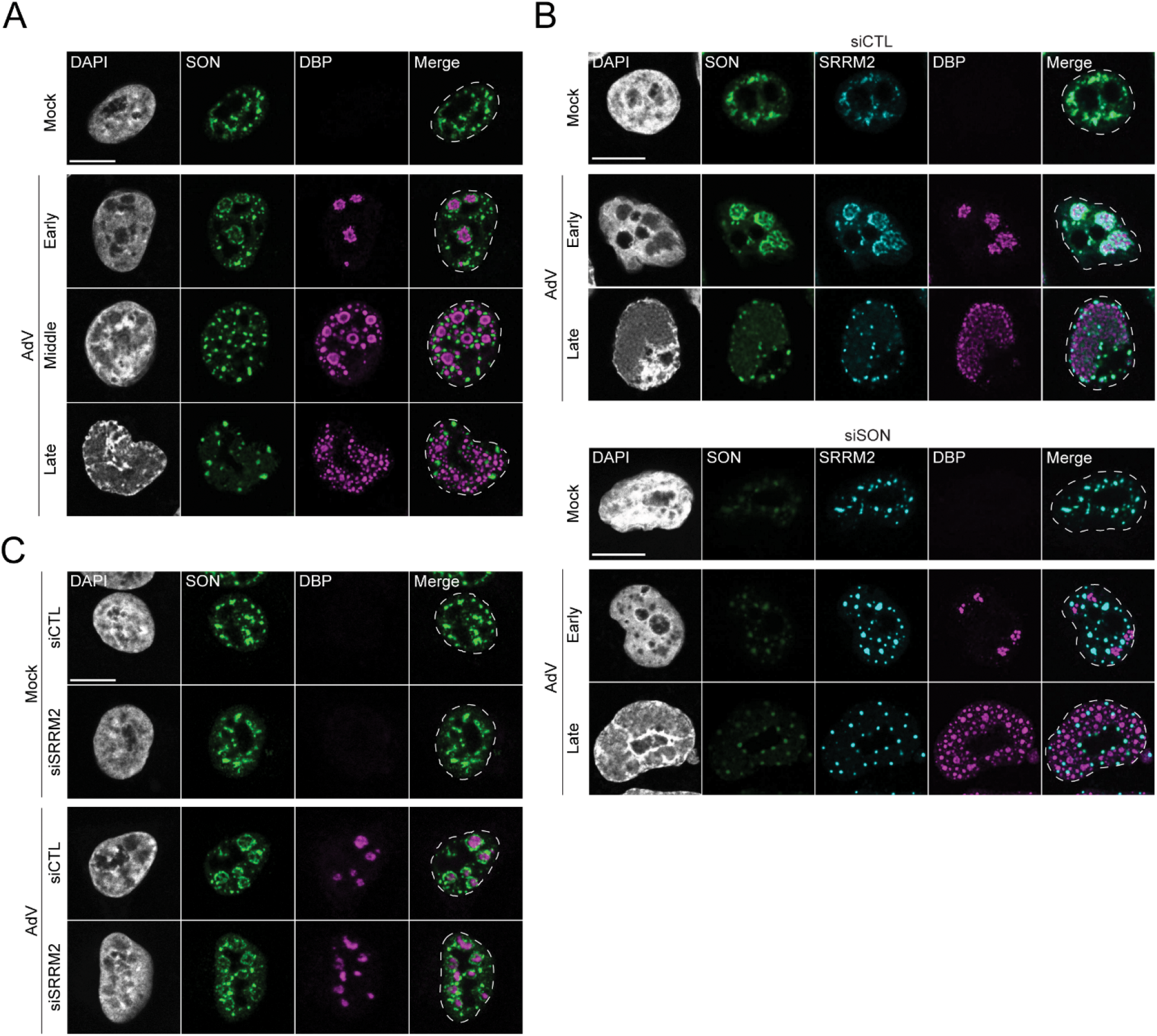
Additional analyses of NS core protein localization patterns during AdV infection. A) Confocal images depicting SON localization in A549 cells following infection with AdV for 20 h. Representative images of cells in different stages of infection are shown. B) Confocal images depicting SRRM2 localization patterns in HeLa mScarlet3-SRRM2 cells following transfection with siCTL (top) or siSON (bottom), then infection with AdV for 20 h. Representative images of cells in different infection stages are shown. C) Confocal images depicting SON localization in A549 cells following transfection with siCTL or siSRRM2, then infection with AdV for 20 h. A-C) DBP (magenta) marks sites of viral replication centers. The nuclear periphery is shown by a dotted white line as assessed by DAPI staining. Scale bar = 10 µm.

**Supplemental Figure 6:**
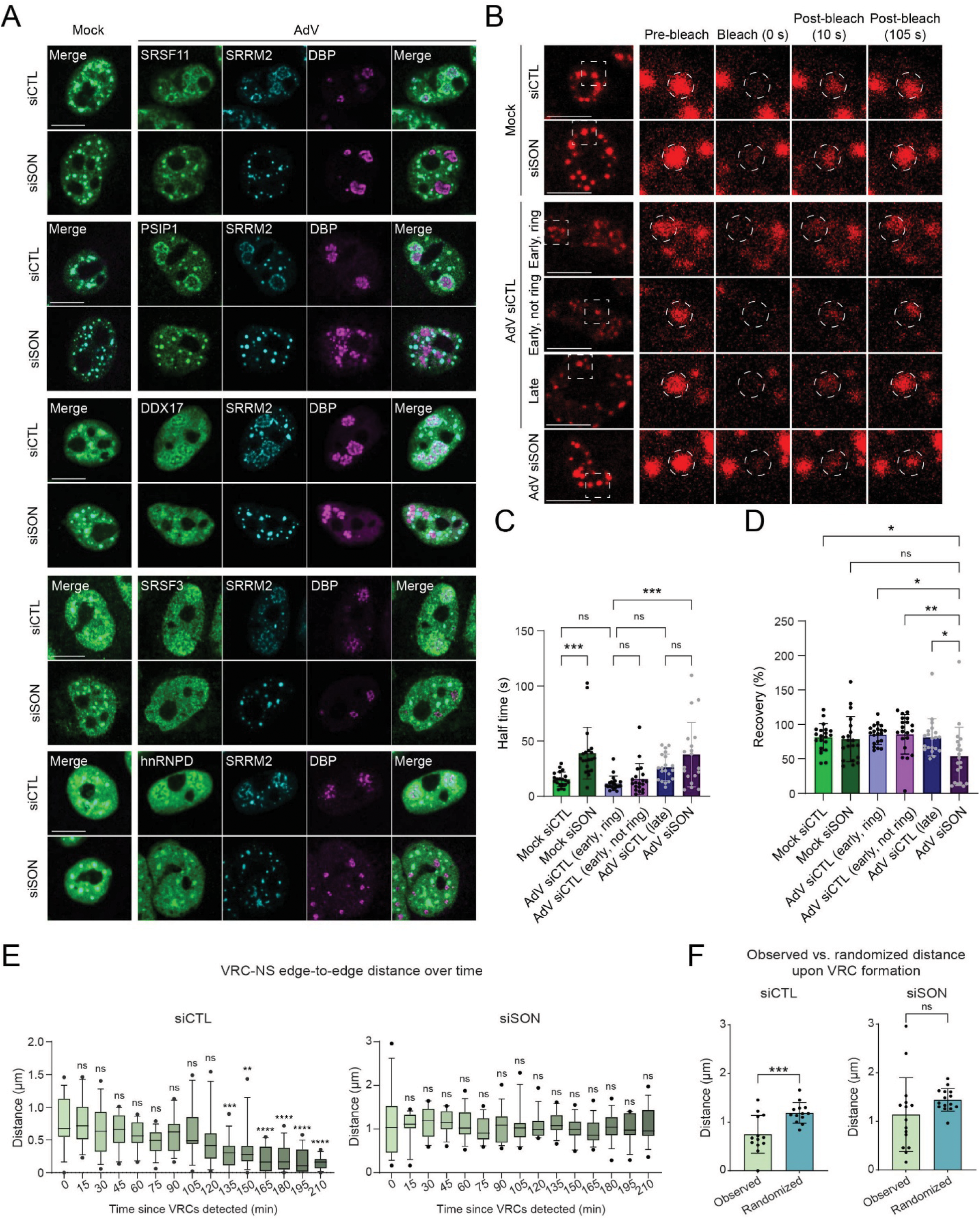
Supporting data corresponding to fluorescence recovery after photobleaching and live-cell analysis of NS. A) Confocal images of proteins in Figure 6 in HeLa mScarlet3-SRRM2 cells relative to viral replication centers. Scale bar = 10 µm. B) Representative live cell images of NS. Scale bar = 10 µm. The dashed white line indicates zoomed area of 6.4 x 6.4 µm. Zoomed area panels show speckles immediately before photobleaching (pre-bleach), immediately after photobleaching (bleach), 10 s after photobleaching, or 105 s after photobleaching. Dashed white circle indicates the bleached area. C) Recovery half times. D) Percentage fluorescence recovery. C-D) Significance was determined by one-way ANOVA with Tukey’s multiple comparisons test. *p≤0.05, **p≤0.01, ***p≤0.001, ****p≤0.0001, ns=not significant. Relevant comparisons are shown. Graphs represent mean +/- standard deviation. E) Average edge-to-edge distances of VRCs and NS at all frames of live-cell imaging for siCTL cells (left) and siSON cells (right). *n* = 13 cells (siCTL) and *n* = 15 cells (siSON). Significance was determined by one-way ANOVA followed by Dunnett’s multiple comparisons test compared to 0 min, with *p≤0.05, **p≤0.01, ***p≤0.001, ****p≤0.0001, ns=not significant. Data are shown as box-and-whisker plots where the central line represents the median, box edges represent the 25^th^ and 75^th^ percentiles (IQR), and whiskers represent the 10-90^th^ percentiles. F) Observed vs. randomized edge-to-edge distances in siCTL cells (left) and siSON cells (right). Each data point represents the average of all VRC-NS edge-to-edge distances within an individual cell. *n* = 13 cells (siCTL) and *n* = 15 cells (siSON). Significance was determined by a two-tailed paired t-test, with *p≤0.05, **p≤0.01, ***p≤0.001, ****p≤0.0001, ns=not significant. Graphs represent mean +/- standard deviation.

